# Cell patterning by secretion-induced plasma membrane flows

**DOI:** 10.1101/2020.12.18.423457

**Authors:** Veneta Gerganova, Iker Lamas, David M. Rutkowski, Aleksandar Vještica, Daniela Gallo Castro, Vincent Vincenzetti, Dimitrios Vavylonis, Sophie G Martin

**Author notes:** These authors contributed equally to this work (Veneta Gerganova, Iker Lamas, David M. Rutkowski).

## Abstract

Cells self-organize using reaction-diffusion and fluid-flow principles. Whether bulk membrane flows contribute to cell patterning has not been established. Here, using mathematical modelling, optogenetics and synthetic probes, we show that polarized exocytosis causes lateral membrane flows away from regions of membrane insertion. Plasma membrane-associated proteins with sufficiently low diffusion and/or detachment rates couple to the flows and deplete from areas of exocytosis. In rod-shaped fission yeast cells, zones of Cdc42 GTPase activity driving polarized exocytosis are limited by GTPase activating proteins (GAPs). We show that membrane flows pattern the GAP Rga4 distribution and coupling of a synthetic GAP to membrane flows is sufficient to establish the rod shape. Thus, membrane flows induced by Cdc42-dependent exocytosis form a negative feedback restricting the zone of Cdc42 activity.

**One Sentence Summary:** Exocytosis causes bulk membrane flows that drag associated proteins and form a negative feedback restricting the exocytic site.

## Main Text

Cells are highly polarized entities that exhibit one (or several) fronts with distinct lipid and protein plasma membrane composition. Reaction-diffusion systems have come a long way in explaining polarity establishment through positive and negative feedback regulations. For instance, in yeast cells, scaffold-mediated positive amplification of Cdc42 GTPase activity underlies the formation of one (or two) cell fronts (*1–4*), with negative feedback promoting oscillations (*5–7*). Physical processes such as cytoskeletal-driven cortical and cytosolic fluid flows, causing advective protein movements, also play important roles in cell polarization (*8*), and can in principle promote polarity establishment (*9*). Membrane flows are predicted to occur and equilibrate membrane tension between spatially segregated zones of exo- and endocytosis, such as in tip-growth of walled pollen tubes (*10, 11*) or in cell migration, as initially proposed in (*12*). However, even though membrane flows have been observed in migrating amoebae, apicomplexans, amoeboid macrophages and neuronal growth cones (*13–16*), their role in cell patterning has not been explored.

We recently described the use of the CRY2-CIB1 optogenetic system in fission yeast (*2*), where blue light activates the cytosolic photosensor domain CRY2PHR (CRY2) to bind its ligand CIBN, linked to the plasma membrane (PM)-interacting domain of the mammalian protein Rit (RitC). These cells grow by tip extension at one or both cell poles, which are sites of endocytosis and exocytosis promoted by local Cdc42 GTPase activity. While in the dark CIBN-RitC is nearly uniform at the PM and blue light initially caused uniform CRY2 recruitment, within 5 min both proteins cleared from most interphase cell poles (Fig 1A, Movie 1). Similarly, in pre-divisional cells, CRY2 and CIBN cleared from mid-cell, where exocytosis is targeted, but re-appeared at non-growing cell poles (Fig S1A, Movie 2). CRY2 depletion after 5 min correlated with local levels of Cdc42-GTP, marked by CRIB-3GFP (Fig 1A-B), the secretory vesicle associated Rab11-like GTPase Ypt3 (Fig 1B, S1B), exocyst subunits Exo70 and Exo84 (Fig S1C-D), and with fast depletion rates (Fig S1E). Thus, light-induced changes in CRY2-CIBN interactions provoke their clearance from zones of exocytosis.

**Figure 1.**
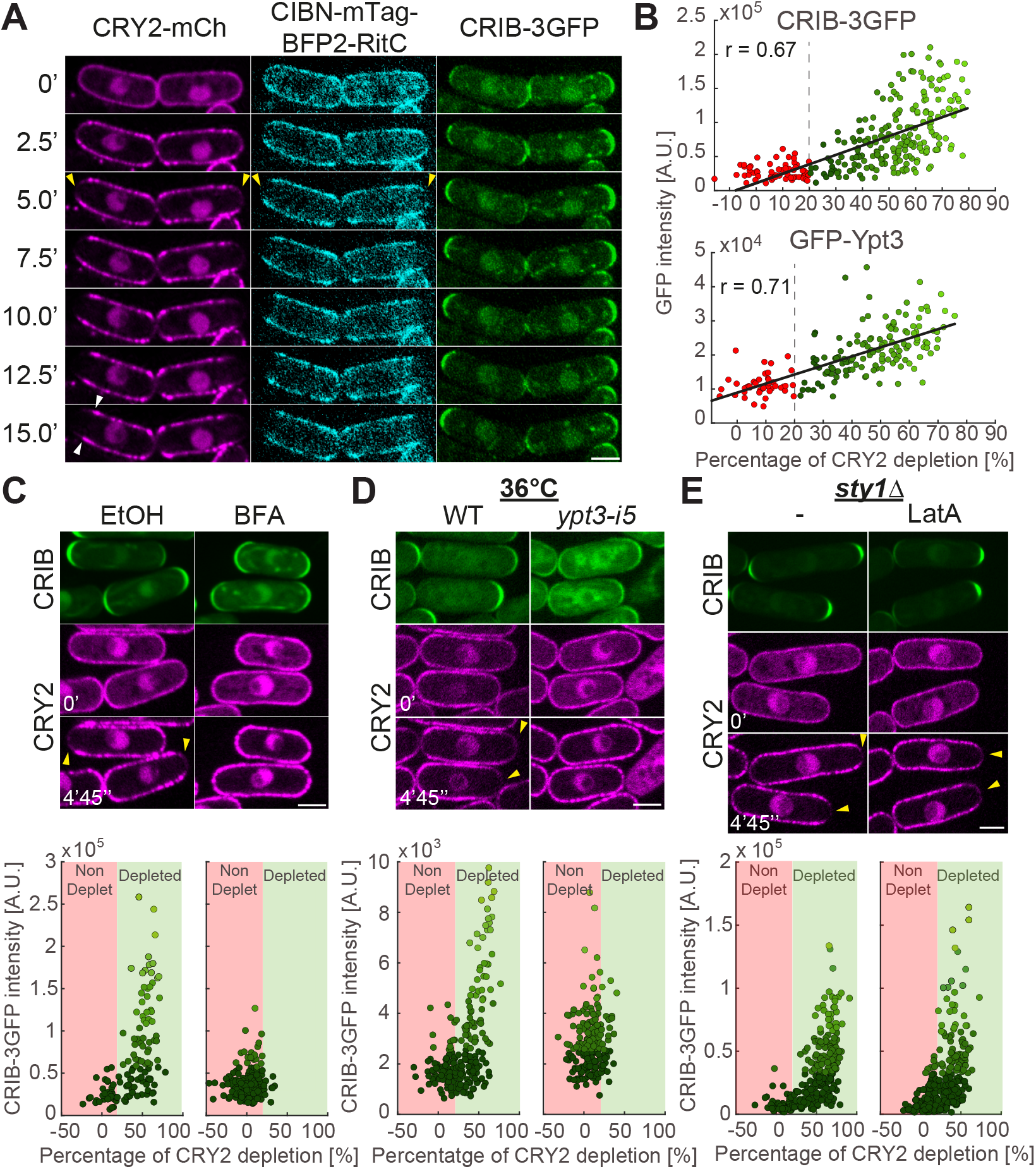
Depletion of membrane-associated CRY2 around sites of secretion. **A.** Time-lapse of CRY2-mCherry, CIBN-mTag-BFP2-RitC and CRIB-3GFP cells grown in the dark. Time 0 is the first timepoint after illumination. Yellow arrowheads indicate depletion zones. White arrowheads point to lateral peaks. **B.** Correlation plot of CRY2 depletion after 5 min and pole GFP intensity of CRIB-3GFP and GFP-Ypt3. Depletion cut-off is indicated by red to green colour change. **C-E.** Cells as in (A) treated with 300μM brefeldin A (BFA) or solvent (EtOH) (C), carrying or not *ypt3-i5* grown at 36°C for 30 min (D) or carrying a *sty1* deletion (*sty1*Δ) treated or not with 50μM Latrunculin A (LatA) (E). The GFP channel is a 5-min temporal sum projection. mCherry shows individual time points. Bottom graphs are as in (B). Data points are coloured with different shades of green according to GFP fluorescence. Scale bars are 3μm in all figures.

CRY2 was not depleted from cell poles upon Brefeldin A (BFA) treatment, which completely stops secretion (*17, 18*), with some cell poles even gaining in CRY2 signal (Fig 1C, S1F-G). CRY2 was also not depleted upon secretion inactivation in the *ypt3-i5* mutant (Fig 1D, S1H). We note that both conditions also caused reduction in CRIB-GFP, perhaps due to loss of positive feedback between growth and polarity (*19*). GFP-Ypt3 intensity was also slightly reduced in BFA-treated cells (Fig S1G). Nevertheless, comparison of cell poles with similar GFP intensity clearly showed that CRY2 does not deplete upon secretion block. Cell poles are also sites of endocytosis, which is strictly dependent on actin patch assembly by the Arp2/3 complex. To test the role of endocytosis, we disrupted all actin structures with Latrunculin A (LatA) in cells lacking the MAPK Sty1, since LatA treatment of WT cells causes MAPK stress signalling and loss of cell polarity (*20*). In LatA-treated *sty1*Δ cells, actin patches were lost, but CRY2 still depleted from cell poles (Fig 1E, S1I). We conclude that CRY2-CIBN depletion can occur by exocytosis alone, but not by endocytosis.

We performed computer simulations to test the hypothesis that deposition and retrieval of membrane material through exo- and endocytic events generates flows strong enough to deplete membrane-associated proteins at growth sites. We modelled proteins as particles with association/dissociation rate constants *k_on_* and *k_off_*, diffusing on a sphere approximating the cell tip, and displaced by exo/endocytotic events over zones of width *σ_exo_* and *σ_endo_* (Fig 2A, Movie 3; see Methods section for details). Particles are displaced radially around the site of vesicle delivery/internalization (Fig 2B), as should occur for proteins in fluid membrane components, flowing to accommodate the change of membrane area on a background of membrane tension. To calibrate the model, we measured rates of fission yeast endocytosis (Table S1).

**Figure 2.**
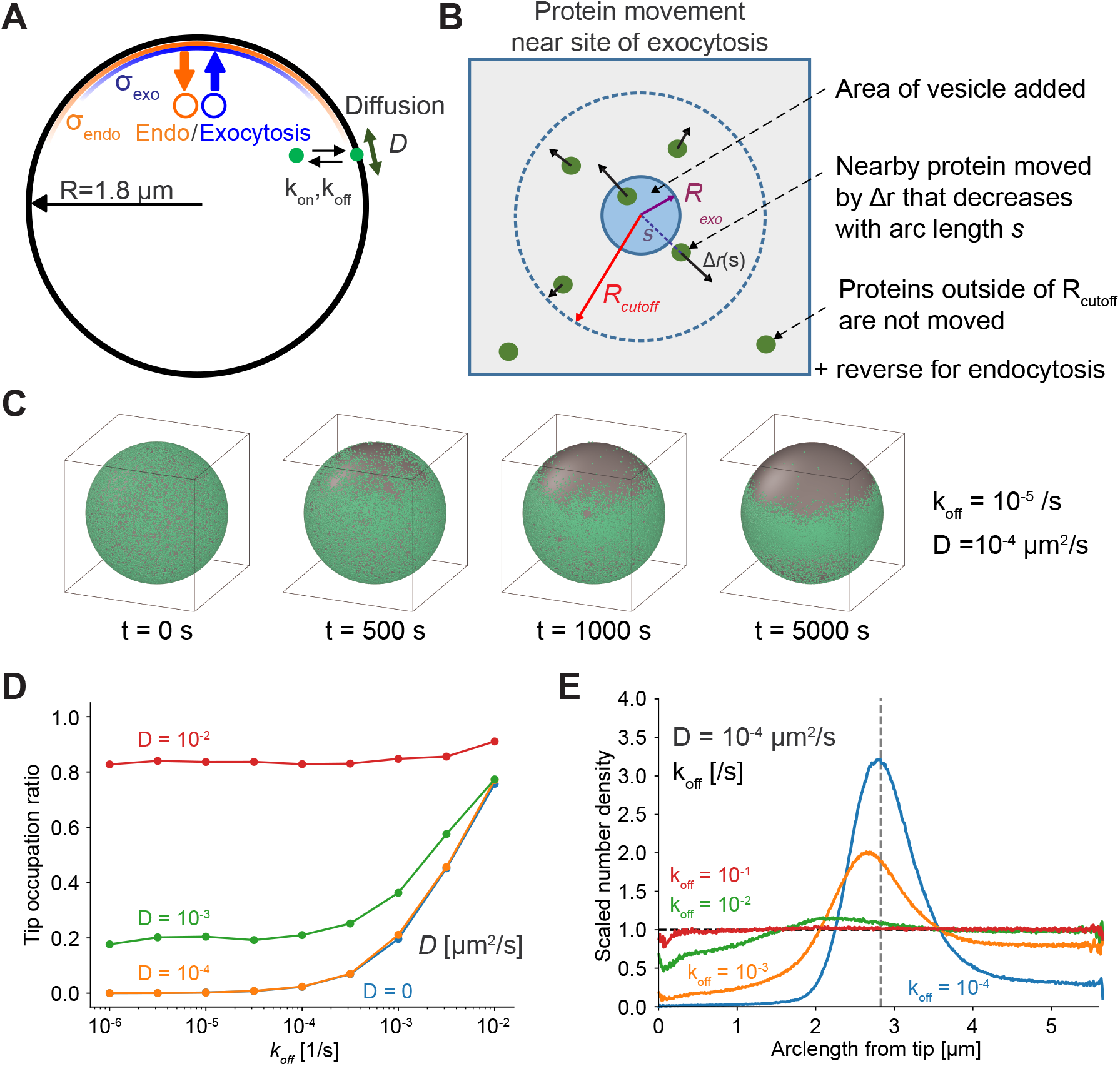
Simulations of membrane-flow-induced concentration gradients. **A.** Particles representing proteins have diffusion coefficient *D* on a spherical cell and exchange with a uniform internal pool with rates *k_off_, k_on_*. Exocytosis (endocytosis) events occur with a distribution of width *σ_exo_* (*σ_endo_*) around a point representing cell tip. **B.** Particle displacement away/towards site of exocytosis/endocytosis. **C.** Depletion of particles starting from uniform distribution occurs for *σ_exo_* < *σ_endo_*, under static conditions (exocytosis rate = measured endocytosis rate). **D.** Tip occupation ratio vs *k_off_* and *D*, same conditions as (C). **E.** Scaled number density vs arclength shows depletion and lateral peak near sphere equator at low *k_off_* values.

Considering conditions where overall membrane delivery by exocytosis balances internalization by endocytosis, and assuming that neither exocytic nor endocytic vesicles carry membrane-associated proteins, we found tip depletion when exocytosis is narrower than exocytosis, *σ_exo_* < *σ_endo_* (Fig 2C, Fig S2A). This is a condition which we verified in cells (Fig S3 and Methods). The active diffusion generated by the randomness of endo- and exocytic events was not by itself sufficient to drive significant concentration gradients when *σ_exo_* = *σ_endo_* (Fig S2A, S2C). Under net growth conditions, where exocytosis exceeds endocytosis by the amount of extra membrane needed for cell growth, depletion was stronger and faster, and occurred even when *σ_exo_* = *σ_endo_* (Fig S2A-C). Net growth conditions in absence of endocytosis (mimicking growth upon LatA treatment) also led to depletion (Fig S2E). Thus, a membrane flow-based model can reproduce the observed depletion from cell poles.

Using the measured endocytosis and exocytosis profiles in our model, we plotted the tip occupation ratio, a measure of depletion, as a function of *k_off_* and diffusion coefficient *D* (Fig 2D). The model makes two key, testable predictions: First, significant tip depletion requires both parameters to be sufficiently small – this is the limit where particle movement and exchange are too slow to counteract flow away from the tip (Movies 3, 4). Second, membrane material flowing away from the cell tip accumulates in a region of enhanced density near the end of the depletion area (Fig 2E). Lateral peak formation requires slow diffusion rates (Fig S2D). Both tip depletion and lateral build-up were stronger in conditions of net growth (Fig S2B).

The finding that depletion is only observed for proteins with slow dynamics (low *k_off_* or *D*) provides an explanation for the acute change in CIBN localization upon light-induced CRY2 interaction. Indeed, light activation promotes not only CIBN-CRY2 binding but also formation of CRY2 oligomers through a distinct binding interface (*21, 22*). CRY2 oligomers form bright foci (see Fig 1A), predicted to reduce dynamics through clustering of membrane anchors. Using FRAP to estimate *D* and *k_off_* (Fig S4), we found that CRY2 interaction causes a >10-fold decrease in CIBN-RitC *D* and *k_off_*, to values consistent with tip depletion in the simulations (Fig 3A). CRY2-CIBN tip depletion was more marked in experiments than predicted by the measured *D* and *k_off_* values, which we reasoned is due to CRY2 oligomerization not only decreasing membrane-unbinding dynamics but also biasing CRY2 binding from the cytosol. This can be accounted for in the model by increasing the rate of particle dimerization on the membrane, which leads to exacerbated tip depletion (Fig S5). We propose that tip depletion events revealed by optogenetics are due to acute clustering-mediated changes in membranebinding properties.

**Figure 3.**
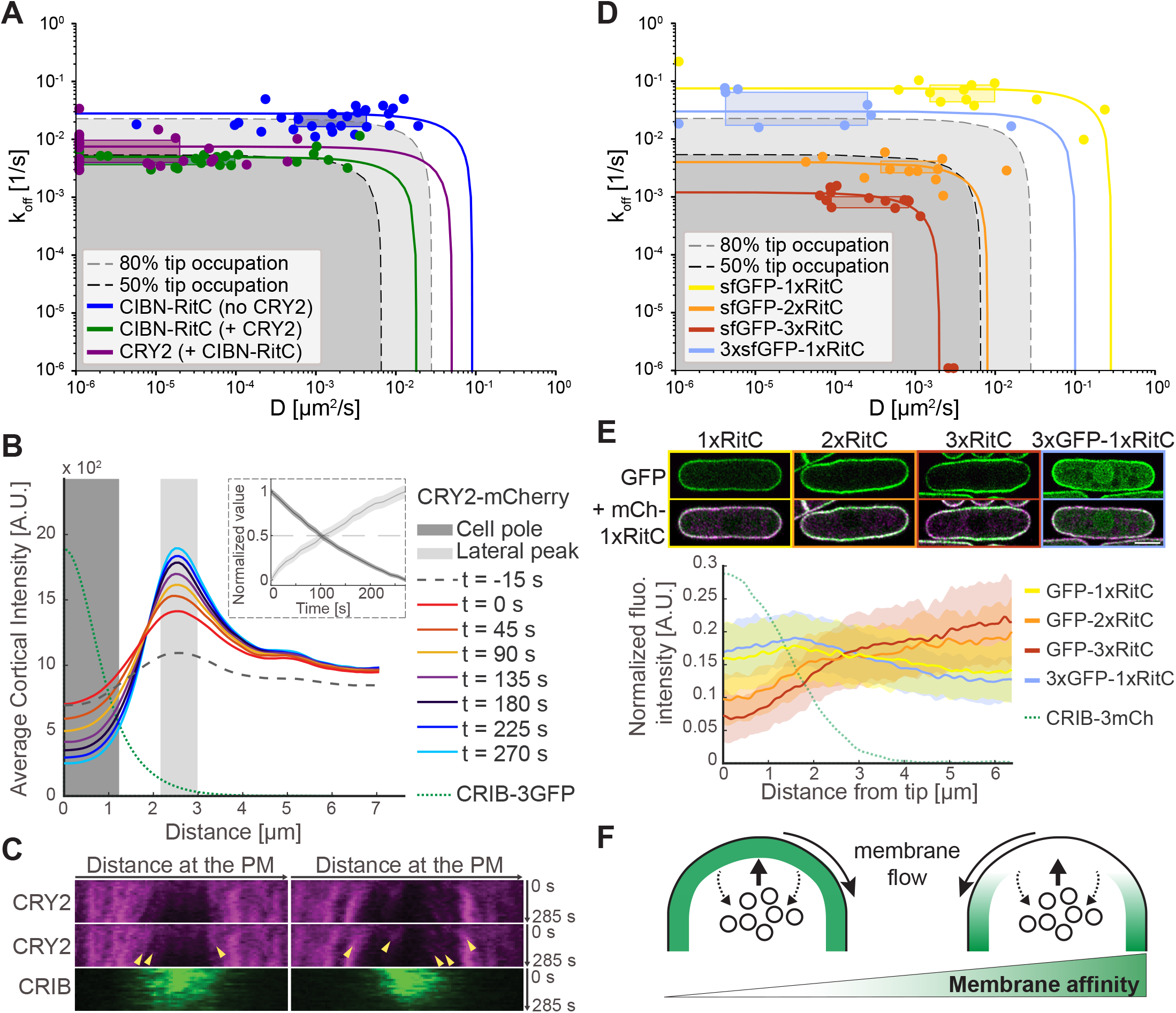
Experimental validation of membrane flows and affinity-dependent protein depletion. **A.** *D* and *k_off_* of CIBN-RitC and CRY2 with or without CRY2 co-expression, derived from FRAP measurements. Points are best *D* and *k_off_* fits for individual cells. Boxes are data point lower and upper quartiles. Lines show *D* and *k_off_* combinations resulting in same overall FRAP recovery (from ridge of averaged R^2^ data fits, Fig S4). Curved corner region corresponds to comparable contributions of *D* and *k_off_*; outer limits to recovery dominated by diffusion or exchange. Grey shaded regions have predicted tip occupation below indicated levels (from growth condition simulations). **B.** CRY2 cortical profiles in time-lapse imaging as in Fig 1A. Time 0 is the first post-illumination image. Please compare to Fig 2E. The green dashed line shows the average CRIB distribution. The inset shows the average fluorescence value over time in the shaded areas. **C.** Kymographs of the cell pole in cells as in Fig 1A. **D.** *D* and *k_off_* of indicated RitC constructs derived from FRAP measurements. Formatting is as in (B). **E.** Localization of indicated sfGFP-tagged RitC constructs in cells co-expressing mCherry-RitC. The graph shows sfGFP-RitC constructs and CRIB signal distribution (from cells as in Fig S6). **F.** Scheme of secretion-induced membrane flow, leading to associated protein depletion according to membrane affinity.

Simulations of protein removal by endocytic vesicles neglecting membrane flows, which were previously shown to maintain the polarization of slow-diffusing transmembrane proteins delivered by secretory vesicles (*23*), generate tip depletion for similar, low values of *k_off_* and *D*. However, in contrast to membrane flow simulations, they do not produce a lateral accumulation peak (Fig S2F-G). Thus, protein accumulation on the edges of the depletion zone is a characteristic feature of the tip depletion model. Concordant with the flow model, a lateral CRY2 peak corresponding to the edge of the CRIB-labelled depletion zone increased over time (Fig 3B, Fig 1A white arrowhead). The rates of CRY2 levels decrease from cell pole and increase at edges were indistinguishable, suggesting they reflect the same molecular process. Kymographs of CRY2 oligomers at cell tips also directly showed events of flow away from the cell pole (Fig 3C). These observations support the existence of bulk membrane flows carrying associated proteins away from sites of polarized secretion.

To test the role of membrane binding in flow-mediated protein patterning without the amplifying effect of oligomerization, we constructed a series of one, two or three tandem copies of the membrane anchor RitC tagged with sfGFP. Estimated *k_off_* and *D* rates extracted from FRAP experiments were higher for 1xRitC than 2xRitC and 3xRitC (Fig 3D). A control 3xsfGFP-1xRitC construct showed only slightly reduced *k_off_* compared to sfGFP-1xRitC, suggesting a small effect of protein size or perhaps weak sfGFP-mediated oligomerization. The measured dynamic parameters of 2xRitC and 3xRitC, but not 1xRitC and 3xsfGFP-1xRitC, fall within the range predicted in the model to cause local depletion (Fig 3D). Indeed, cell pole depletion was observed for 2xRitC and more prominently 3xRitC, but not 1xRitC constructs. This is particularly visible in comparison with the uniform localization mCherry-1xRitC co-expressed in these cells (Fig 3E, Movie 5). Signal depletion at cell poles correlated with Cdc42-GTP levels (Fig S6). Thus, membrane affinity is a strong predictor of protein distribution around sites of polarized exocytosis (Fig 3F). The dependence of protein distribution on membrane affinity should invite careful consideration of this parameter when designing probes to evaluate lipid distribution.

To probe the role of membrane flows in cell patterning, we focused on the Cdc42 GTPase activating protein (GAP) Rga4, which is excluded from cell poles and forms membrane-associated clusters distributed in a ‘corset’ pattern, reminiscent of the lateral peaks described above (Fig 4A) (*24, 25*). Previous work suggested that cell side-localized Cdc42 GAPs control cell width by restricting the size of the Cdc42 active zone (*24, 26, 27*). Extensive structurefunction analysis of GFP-tagged Rga4 fragments, described in Fig S7 and supplemental text, indicated oligomerization by coiled-coils (cc) and the presence of a cortex-binding domain (CBD) comprising two membrane-binding motifs. A minimal cc-CBD Rga4 fragment recapitulated the main features of Rga4 localization, forming clusters partly depleted from cell poles (Fig 4B) and was used to probe how a simplified endogenous protein may be affected by membrane flows. Truncating the coiled coils led to faster membrane-binding dynamics and repopulation of the cell poles (CBD only, Fig 4B-D). Tandem multimerization of the CBD (4xCBD) increased membrane affinity and restored depletion from cell poles (Fig 4B-D), as did artificial CBD oligomerization by fusion with CRY2 (Fig S8A). Truncating one of the membrane-binding motifs (cc-CBD-Δ2) to reduce membrane affinity led to faster protein dynamics, as confirmed by FRAP analysis (Fig 4D) and TIRF microscopy (Fig S8B-C, Movie 6-7). This truncation caused enrichment at cell poles (Fig 4B-D), indicating affinity of the coiled coil region for cell poles. Increasing membrane-affinity by multimerization (4x cc-CBD-Δ2) restored tip depletion and enrichment to cell sides (Fig 4B-D). Thus, even with a preferential association at cell poles, the oligomerization and two membrane-binding motifs of Rga4 ensure a sufficiently long membrane residence time that couples Rga4 to membrane flows towards cell sides.

**Figure 4.**
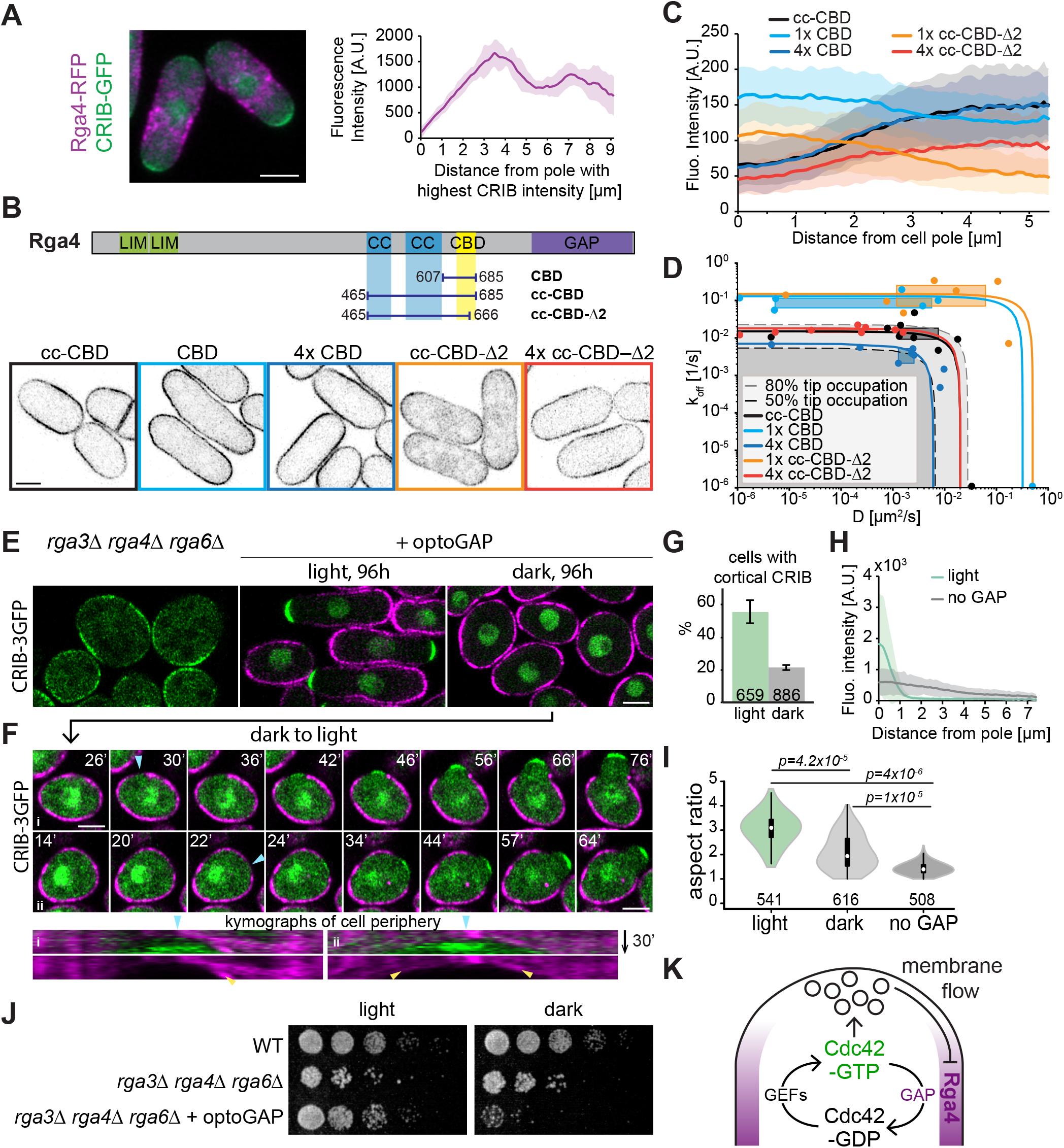
Membrane flow-induced Cdc42 GAP depletion forms a negative feedback restricting the secretion zone. **A.** Maximum projection image of Rga4-RFP and CRIB-3GFP. The graph shows average and stdev of Rga4 distribution along cell length in 11 cells. **B.** Scheme and localization of the minimal cc-CBD Rga4 fragment, as well as single or 4x tandem CBD and cc-CBD-Δ2 constructs. **C.** Average cortical distribution of indicated constructs. Shaded areas show standard deviation. **D.** Diffusion and membrane detachment rates of Rga4 fragments as in (B) derived from FRAP measurements. Formatting as in Fig 3A. **E.** *rga3*Δ *rga4*Δ *rga6*Δ *CRIB-3GFP* cells expressing or not optoGAP continuously grown in the light or dark for 96h. **F.** Cells as in (E) grown in the dark and shifted to light at time 0. Arrowheads indicate optoGAP depletion. Kymographs show lateral movement of OptoGAP. **G.** Percentage of optoGAP-expressing cells as in (E) with local CRIB cortical signal. **H.** CRIB-GFP cortical profiles in light-grown optoGAP cells vs. cells not expressing optoGAP as in (E). **I.** Aspect ratio of cells as in (E). **J.** Serial dilution of indicated cells on rich medium. **K.** A double negative feedback regulates the size of the Cdc42-GTP zone. Cdc42-GTP promotes secretion, which induces membrane-associated flow of Rga4 GAP to cell sides where it restricts Cdc42 activation, thus constraining the Cdc42-GTP zone.

To directly test the importance of the GAP lateral localization, we constructed a synthetic GAP (optoGAP) consisting of CRY2, the membrane-binding RitC, and the Rga4 GAP domain. OptoGAP is predicted to decorate the plasma membrane uniformly in the dark, but couple to membrane flows and deplete from cell poles in the light, due to CRY2 oligomerization. When expressed in the light in cells lacking all endogenous Cdc42 GAPs, which exhibit poor growth and a round morphology with very large zones of Cdc42 activity (*28*), not only was optoGAP depleted from zones of Cdc42 activity (Fig 4E), but the zone size was strongly reduced (Fig 4E, 4H), restoring viability and the rod cell shape (Fig 4I-J). Upon growth in the dark, optoGAP was present around the cell periphery (Fig 4E) and was toxic to the cells (Fig 4J), the majority of which showed no cortical CRIB (Fig 4G) and regained a round morphology (Fig 4I). Further return of these cells to the light led to acute local optoGAP depletion coinciding with a zone of Cdc42 activity, with lateral flow movements captured in kymographs, and outgrowth of a tube of restricted width (Fig 4F, Movie 8). We conclude that coupling of Cdc42 inhibition to membrane flows, which are themselves induced by Cdc42 activity, forms a spatial negative feedback sufficient to restrict zones of Cdc42 activity and define cell width (Fig 4K).

Because membrane flows are a consequence of polarized secretion, they are likely to be prevalent in cells and pattern membrane-associated proteins, coordinating with the cytoskeleton that can independently tune membrane rheology (*29, 30*). Besides Rga4, membrane flows may promote the lateral localization of several proteins that organize the yeast cell shape and division, and explain their exclusion from both cell poles in wildtype cells, but from the only growing pole in monopolar mutants (for instance Rga6, Mid1, Blt1, Cdr2, Skb1 (*25, 27, 31–35*)). Membrane flows may also help couple endocytosis around sites of exocytosis, which also promote polarity establishment (*36, 37*). Other proteins localize to secretion sites despite membrane flows, thanks to fast membrane unbinding, directed cytoskeletal transport or preferred affinity to static mobile components. For instance, pole-enriched Pom1 shows association to microtubule-transported factors and very fast membrane unbinding dynamics (*38, 39*). The Cdc42 GTPase also has to resist dilution from secretion-derived membrane addition (*40*). These principles of membrane-associated protein patterning are likely valid in all eukaryotic cells. Indeed, precise spatial distribution of exo- and endocytosis, predicted to induce membrane flows, accompanies cellular processes ranging from tip-growth in fungi and pollen tubes (*41*), formation of neuronal connections, cilia and immunological synapses (*42, 43*), cell migration to cytokinesis (*44, 45*). Furthermore, because protein localization can be altered by a simple change in membrane unbinding rates, it may be acutely modified through post-translational modifications that modulate membrane affinity and/or oligomerization.

## Acknowledgments

We thank Stephan Gruber (Unil), Jan-Willem Veening (Unil), Serge Pelet (Unil), Laura Merlini (Unil), and Olivia Muriel-Lopez (Unil) for careful reading of the manuscript.

## Author contributions

IL, VG, AV and SGM conceived the project. IL performed all experiments in Fig 1, 3B-C, S1 and S3. VG performed all other experiments in Fig 3, 4 and S6-8. DMR and DV conceptualized the simulations and FRAP analysis methods. DMR performed all simulations and FRAP analyses. AV performed the initial RitC multimerization experiments. DG performed initial Rga4 structure-function analysis. VV provided technical support. SGM and DV coordinated the project and acquired funding. SGM and DV wrote the first draft of the manuscript, which was revised by all authors.

## Data and materials availability

All data and reagents are available upon reasonable request to the corresponding authors.

## Supplementary Materials

### Supplemental Text

The text below describes the structure-function analysis of Rga4 fragments presented in Fig S7.

#### Rga4 structure-function analysis

Through structure-function analysis of GFP-tagged Rga4 fragments, we defined determinants of membrane association and cluster formation. Consistent with previous observations (*25*), a cortex-binding domain (CBD; aa 623-685) was both necessary and sufficient for membrane association (Fig S7A-B). This region could be replaced by RitC to target Rga4 N-terminus to the cortex. CBD sequence analysis revealed two conserved sequences: a weak amphipathic helix prediction (motif 1) and a region rich in basic residues, predicted by a high BH score (*46*), (motif 2) (Fig S7A). Mutation of key residues in these two motifs did not individually block cortical binding, but together resulted in cytosolic localization. This was the case whether the mutations were introduced in a short Rga4 fragment containing coiled-coils (cc) and the CBD (cc-CBD), or whether they were introduced in the full-length Rga4 expressed from the native genomic locus (Fig S7C). Full-length Rga4 containing mutations in both membrane-association motifs, which was cytosolic, was non-functional, as shown by the round shape of cells also lacking the other two Cdc42 GAPs Rga3 and Rga6 (Fig S7D). Thus, these data identify two membrane-binding motifs in the CBD, which are together essential for Rga4 localization at the plasma membrane.

We noted that full-length Rga4 with these mutations formed cytosolic clusters (Fig S7C), as previously observed upon overexpression of cytosolic Rga4 (*26*). The N-terminal fragment lacking the CBD also formed cytosolic condensates (Fig S7B), which disappeared upon exposure to high temperature (37°C for 6h) or 5% 1,6-hexanediol, thought to interfere with weak hydrophobic interactions (Fig S7E), suggesting phase-separation properties. Full-length Rga4 cytosolic clusters were also lost under these conditions (Fig S7F). Importantly, condensates were absent upon further truncation of the coiled coils (Fig S7B). The Rga4 N-terminus containing the LIM domains likely also contribute to condensate formation, as we did not observe condensates in cytosolic fragments lacking this region (Fig S7C, fragment 9). We conclude that Rga4 forms larger molecular assemblies for which the coiled coils and perhaps other determinants are necessary.

As described in the main text, a minimal Rga4 fragment containing both coiled coils and CBD (cc-CBD) recapitulated the main features of Rga4 localization, as it formed membrane-associated clusters that were partly depleted from cell poles (Fig 4B-C, Fig S7C; see also Fig S8C). We note here that the relocalization and enrichment at cell poles observed upon truncation of the second membrane-binding motif (Fig 4B-C) also occurred upon mutations in motifs 1 and 2 (Fig S7C; fragments 7 and 8). We hypothesize that the reported interaction of the coiled-coil region with Pom1 kinase may bias its binding to the cell poles region where Pom1 accumulates (*25, 39*).

### Materials and Methods

#### MODEL METHODS

##### Simulations of effects of exocytosis and endocytosis on protein distributions

To investigate conditions that lead to depletion of proteins from a growing cell tip, we simulate the motion of non-interacting particles moving along a surface representing the plasma membrane under the effects of membrane binding/unbinding, diffusion, and membrane delivery/removal by exocytosis/endocytosis. The biophysical assumptions of our model are presented together with the model implementation below. For simplicity, the particles in these simulations are restricted to the surface of a sphere with radius 1.8 μm with a fixed point on the sphere surface specified to be the growing tip. All arc lengths are measured with respect to this fixed point.

###### Time Step

The simulation was evolved through time primarily by using the Gillespie algorithm (*47*). An event is chosen from the list of possible events that include particle association and dissociation from the membrane, endocytosis, and exocytosis (excluding diffusion). The probability of a specific event occurring is given by the ratio of the event rate to the sum of the rates of all possible events (i.e. *P_i_* = *r_i_*/∑η*r_j_* where *r_i_* is the rate of event *i*). The time until the selected event occurs is calculated as Δ*t* = –ln(*u*′) /∑*r_j_* where *u*′ is a uniform random number between 0 and 1.

###### Diffusion

Particle diffusion, with diffusion coefficient *D*, is implemented with Brownian dynamics. The magnitude of the displacement due to diffusion over time Δ*t* is calculated as 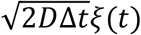 independently in both directions in the plane tangent to the sphere, and *ξ*(*t*) is a random Gaussian distributed variable with zero mean and unit variance. After calculating the displacement in the tangent plane, the positions of each particle are then projected down to the sphere surface while keeping the distance conserved to correctly describe diffusion on a spherical surface (*48*). Since particles are assumed to diffuse independently of each other, we only calculated the displacement due to diffusion prior to endocytosis or exocytosis events, or before saving the data (except for a model with particle dimerization described below). We ensured that this implementation of diffusion on the sphere surface reproduced the correct scaling behaviour for the angular diffusion (*48*) by measuring the change in the polar angle, 〈*θ*(*t*〉^2^), as a function of time up to 1000 s as shown in Fig S9A.

###### Membrane binding and unbinding

Particles on the sphere undergo an unbinding event with rate *k_off_* where they are removed from the sphere surface and placed in an internal pool representing the cytoplasm. Particles in this internal pool are placed back on the surface anywhere with uniform probability with rate *k_on_*. This process assumes a sufficiently fast cytoplasmic diffusion to ensure uniform cytoplasmic concentration. We fix *k_on_* = 0.01 s^−1^ since within our independent particle approximation this rate should only affect the average number of particles on the surface but not the relative level of tip depletion at steady state. To calculate smooth concentration profiles, there are on average approximately 50,000 particles on the surface at steady state in the simulations.

###### Particle movement by exocytosis and endocytosis

We assume that exocytosis and endocytosis near cell tips create a local flow that displaces lipids and proteins around the site of the vesicle event (*40, 49*). Our assumption is supported by the appearance of a flat plasma membrane in electron micrographs of fission yeast cell at cell sides and tips, except at sites of vesicle traffic or around eisosomes (*50–52*), similar to *S. cerevisiae* (*52, 53*). This observation indicates the presence of a membrane tension, which, combined with turgor pressure, can flatten the vesicle membrane delivered by exocytosis (*54, 55*), causing lateral flow. Similarly, flow of lipids towards endocytic vesicles must occur over a sufficiently wide area, preventing membrane strains larger than a few % that would otherwise lead to formation of membrane pores (*56*).

As many of the details of cellular membrane hydrodynamics remain unresolved (*57*), here we adopt a simplified description. We simulate the movement associated with exocytosis as occurring instantaneously (i.e. no diffusion or membrane binding/unbinding occurs during the movement of the particles for either exocytosis or endocytosis). Yeast membranes contain nearly immobile transmembrane proteins, likely connected to the cell wall (*58*). Thus, similar to animal cells, we anticipate a fraction of the membrane, as high as 50%, to be immobile, with the fluid component undergoing Darcy-type flow (*57, 59, 60*). The fluid component of the membrane could be heterogeneous, containing lipids and proteins with different diffusion coefficients, a parameter that we vary in the model. Peripherally-associated membrane proteins are assumed to be physically and hydrodynamically coupled to this lipid flow. To simulate the effect of slow equilibration of membrane tension over long distances (*30*), we also introduce a cutoff distance to the flow induced by individual exocytosis and endocytosis events. As we show below, our results on tip depletion are robust with respect to the precise values of the mobile fraction and cutoff distance.

In the simulation, exocytosis and endocytosis events take place on the surface of the sphere and affect the positions of nearby particles. The location of these events is selected randomly from an experimentally determined distribution (see below). Particles on the surface at arc length distance *s* from the event are pushed away radially from the center of exocytosis as shown in Fig. 2B by arc length distance Δ*r*(*s*). It is assumed that the difference between the spherical cap surface area corresponding to mobile proteins and lipids, *A_cap_*, measured at the final and initial arc length positions of the particle from the center of exocytosis equals the surface area of the exocytotic vesicle *A_exo_*:

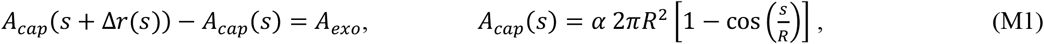

where *s* is the arc length distance, *R* is the radius of the sphere, and parameter *α* describes the fraction of mobile proteins and lipids between 0 (no mobility) and 1 (all mobile). This gives:

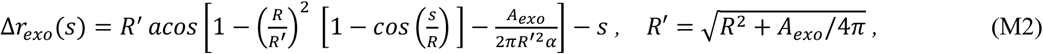

where *R*′ is the radius the sphere would have to accommodate the membrane area contributed by the exocytotic vesicle. The displacement is only applied to particles up to an upper arc length distance, *s* < *R_cutoff_*, beyond which we assume the effects of membrane delivery are not felt immediately. The small difference between *R* and *R*′ was introduced to avoid artificial accumulation of particles at the opposite sphere end in the absence of a cutoff *R_cutoff_* but is otherwise insignificant for the simulations with a cutoff distance smaller than the arc length corresponding to the half-sphere. For an endocytosis event, particles within the cutoff distance *R_cutoff_* are moved radially towards the endocytic point, similar to the calculation for exocytosis, with displacement magnitude:

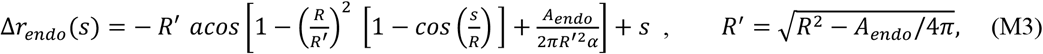

where *A_endo_* is the surface area of the endocytic vesicle. When the displacement magnitude Δ*r_endo_*(*s*) calculated by Equation (M3) is larger than *s*, particles are moved to the endocytosis event point, for the simulations where endocytosis does not contribute to internalization; for the model with endocytic removal they are internalized into the internal pool of particles.

###### Distribution of exocytosis and endocytosis

The distribution for exocytosis/endocytosis is based on the experimental distribution as function of arc-length distance from the cell tip, *P_exp_*(*s*), measured by imaging the profile along a medial confocal plane. The distribution *P_exp_*(*s*) can be well described by a Gaussian of standard deviation *σ_exo_* and *σ_endo_* for exocytosis and endocytosis, respectively. To convert this two dimensional distribution to a three dimensional one, we assume that the experimentally observed profiles account for events within a slab of width on the order of the microscope’s resolution along the axis perpendicular to the glass slide (*b* = 0.71 μm) (Fig S9B). Thus, assuming cylindrical symmetry, the distribution of an event at arc length *s* from the position defined as cell tip used in simulations is calculated as

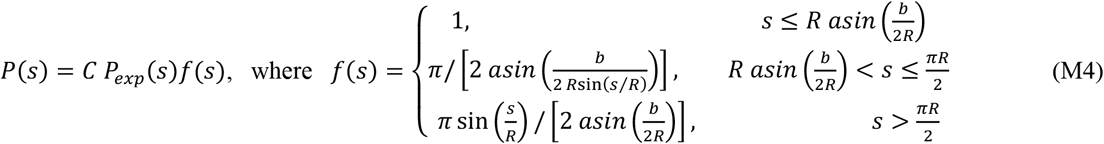

where *P_exp_*(*s*) is the experimental 2D probability distribution, *P*(*s*) is the adjusted 3D probability distribution used in simulations, *C* is a normalization constant and *f*(*s*) is the fraction of the circumference of the circle at a given value of *s* outside the slab region (i.e. thin line region of the circle circumference divided by the thick line region in Fig. S9B). The expression for *f*(*s*) initially assumes that for *s* > *πR*/2 the shape is that of a spherocylinder (as this is the shape of the actual cell) which is then mapped back onto a sphere by keeping the density of events constant by multiplying by sin(*s*/*R*). Agreement between experimental and simulated *P*(*s*) considering only the slab region is shown in Fig. S9C. The azimuthal angle for the exocytotic or endocytic event is picked from a uniform distribution.

###### Rates of membrane area delivery by endocytosis and exocytosis

We calibrated the model based on measurements performed as described in the Experimental Materials and Methods section.

Firstly, the measured average rate of old end tip growth was 0.0305 ± 0.0151 μm/min (n = 75 cells; average ± stdev), which, multiplying by 2*πR* corresponds to net membrane area added of 0.345 μm^2^/min = 20.7 μm^2^/hr.

Secondly, we measured the rates of endocytosis at growing old end tips by counting the number of Fim1 patches in a medial section and extrapolating to the whole cell tip area. Since the Gaussian profile of the endocytosis distribution extends beyond the hemispherical tip region, we considered separately the medial section rates at the hemi-spherical region (arclength from tip 0 to *πR*/2, with 24.35 ± 10.56 events/min) and cylindrical segment (arclength *πR*/2 to *πR*, with 5.26 ± 3.85 events/min counting both sides). The fraction of events in a medial section of the hemi-spherical region would be 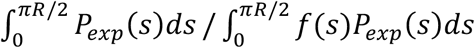 This ratio is 0.233 using the measured *σ_endo_*, leading to 104.53 endocytosis events/min for the whole hemi-spherical region. The fraction of cylindrical area imaged in a medial section is *b*/(*πR*) = 0.126, leading to 41.62 endocytosis events/min in the cylindrical region around the cell tip. Summing the numbers for hemi-spherical and cylindrical regions provides a rate of *r_endo_* =146.2 /min = 2.44 /s as listed on Table S1. This corresponds to endocytosed membrane area of 0.939 μm^2^/min, using the estimate of endocytic vesicle radius in Table S1.

Thirdly, we estimated the rates of exocytosis by assuming that the rate of membrane area delivered by exocytosis is equal to the sum of net area added due to growth plus the membrane area endocytosed (sum = 1.28 μm^2^/min). Using the estimate of exocytic vesicle radius in Table S1 leads to *r_exo_* = 40.9 /min = 0.68 /s as listed on Table S1.

###### Measurement of tip occupancy ratio and reference parameters

The tip occupancy ratio is calculated by dividing the density of particles at an arc length of *πR*/4 = 1.41 μm (a quarter of half sphere total tip arc length) away from the top tip of the sphere by the density of particles in the remaining area of the sphere. The reference parameter values are listed in Table S1, at the bottom of this section.

###### Effect of parameters k_on_, mobile fraction a, and cut-off distance R_cutoff_

Results presented in the main figures and Fig. S2 used values of *k_on_* = 0.01 *s*^−1^, *α* = 0.5, *R_cutoff_* = 2 μm. We also examined the effect of varying these parameters. As expected, the steady state reached is independent of *k_on_* (Fig. S9D). We find that decreasing *α* to a value of 0.1 (low mobile fraction) results in larger depletion and higher lateral peak, while increasing *α* to a value of 1.0 (highest mobile fraction) reverses the trend (Fig S9E). Decreasing the value of *R_cutoff_* to 1.0 μm decreases overall depletion and lateral peak magnitude, shifting the peak towards the left, while increasing *R_cutoff_* to 2.5 μm reverses the trend (Fig S9F).

###### Dimerization simulations

To investigate the effect that oligomerization has on the extent of tip depletion and lateral peak height, we performed additional simulations in which we allow surface particles representing proteins to bind together to form dimers (Fig. S5A). To avoid artificial concentration gradients as a result of implicit energy input to the system, we implemented a scheme designed to satisfy detailed balance in equilibrium, in the absence of exocytosis and endocytosis. In addition to association and dissociation of particles between cytoplasm and surface representing the plasma membrane (reaction 1 in Fig. S5A with rate constants *k_on_, k_off_*), dimers can be formed in two ways: either two surface particles can bind reversibly to form a dimer (reaction 2 in Fig. S5A with rate constants *k_on_*, *k*___) or a particle from the bulk cytoplasm and a surface particle can bind reversibly to form a dimer (reaction 3 in Fig. S5A with rate constants *k*_*on*2_, *k*_*off*,2_). While two particles are bound in the dimer state they have the same position and they move together under diffusion, exocytosis, and endocytosis. To distinguish between the effects of oligomerization versus diffusion, as well as for simplicity, we used the same diffusion coefficient for surface monomers and dimers.

With rate constants in units of inverse time, detailed balance in absence of exocytosis and endocytosis requires 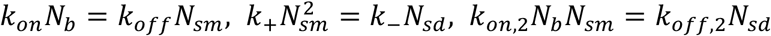, where *N_b_* is the number of bulk particles in the cytoplasm, *N_sm_* is the number of surface bound particles as monomers, and *N_sd_* is the number of surface particles in dimers. We also have *N_b_* + *N_sm_* + *N_sd_ = N_tot_*, where *N_tot_* is the total number of particles which is constant throughout the simulation. Simultaneously solving these four equations results in expressions for the number of particles in each state in equilibrium, *N_b,eq_*, *N_sm,eq_*, and *N_sd,eq_* and a fourth equation, the detailed balance condition, for the reaction rate constants which can be written as *k_on_* = *k*_*on*,2_ *k*_-_ *k_off_* /(*k*_*off*,2_ *k_on_*). In these simulations, we kept *k_off_* = *k*___ = *k*_*off*,2_ and *k_on_* = 0.01 *s*^−1^.

To investigate the effect of dimerization on tip depletion, we then turned on exocytosis and endocytosis. We define a parameter describing the effect of cooperativity, which is the ratio of equilibrium rate of particle association to the surface through dimer formation (reaction 3) versus the equilibrium rate of bulk cytoplasm particle attaching to the surface as monomer (reaction 1): *p* = *k*_*on*,2_*N_b,eq_N_sm,eq_*/(*k*_*on*,3_*N_b,eq_*) = *k*_*on*,2_*N_sm,eq_*/*k_on_* The value of *p* thus implicitly defines the value of rate constant *k*_*on*,2_, as well as that of *k*_+_ through detailed balance.

All reactions, except dimer formation through surface monomers, were handled with the Gillespie algorithm, similar to the case without dimerization described earlier. When a cytoplasmic particle binds to the surface to form a dimer, it is placed at the same position as a randomly-picked surface monomer particle. In the reverse reaction, one of the two particles in the dimer is selected to become a cytoplasmic particle while the other maintains its original position.

To handle the reaction of two surface particles binding together to form a dimer we used the method implemented in Smoldyn (*61*), where particles associate when separated by a distance less that a binding radius, *σ_b_*, and are placed at an unbinding radius, *σ_u_*, upon unbinding into two surface monomers, using *σ_b_* = *σ_u_*. To determine *σ_b_* we followed the 3D reaction scheme of Smoldyn as a guide (where *σ_b_* is determined based on the timestep, diffusion coefficient, and *k*_+_), however as reactions in 2D involve additional complexities (*62, 63*), <u>w</u>e adjusted *σ_b_* by explicitly checking that detailed balance is satisfied in simulations without endocytosis or exocytosis. Values of *σ_b_* ranged from 5·10^-5^ to 0.05 μm as *p* was varied. The association reactions were checked over a fixed time interval Δ*t_fixed_* = 0.01 *s*. Diffusion of surface particles was implemented every Δ*t_fixed_* and prior to any endocytosis or exocytosis event. Dimerization between a pair of particles occurs if they are at a distance less than or equal to *σ_b_* at the end of such a timestep. The unbinding reaction of a dimer into two surface particles is handled by the Gillespie algorithm and the rate constant *k*___ where the particles are placed a distance of *σ_u_* away from each other with random orientation. Due to the additional computational costs associated with calculating distances between particle pairs, we reduced the total number of particles to *N_tot_* = 5000 and simulations were run for 10,000 s.

We find that as *p* increases, the number of dimers at steady state increases at the expense of monomers on the surface and cytoplasm particles (Fig. S5B). The tip depletion and height of lateral peak also increases as a function of *p* for fixed values of *D* and *k_off_*, when these were taken to be close to those of CRY2+CIBN-RitC (Fig. S5C-E). This result shows that dimerization, and more generally, oligomerization, can lead to an enhancement of tip depletion and lateral peak.

**Table S1.** Reference parameter values used in simulations.

| Variable | Value | Description |
| --- | --- | --- |
| $R$ | $1.8 \mu\text{m}$ | Radius of model sphere, approximation to *S. pombe* radius |
| $R_{cutoff}$ | $2 \mu\text{m}$ | Cutoff range for exocytosis/endocytosis effect on sphere surface |
| $\alpha$ | 0.5 | Mobile membrane fraction |
| $R_{exo}$ | $50 \text{ nm}$ | Radius of individual exocytotic vesicles (50, 51, 64) |
| $A_{exo}$ | $3.14 \times 10^{-2} \mu\text{m}^2$ | Surface area of individual exocytotic vesicles |
| $R_{endo}$ | $22.6 \text{ nm}$ | Radius of individual endocytotic vesicles based on measurements in *S. cerevisiae* (53) |
| $A_{endo}$ | $6.42 \times 10^{-3} \mu\text{m}^2$ | Surface area of individual endocytotic vesicles |
| $r_{endo}$ | 2.44/s | Rate of endocytosis events [This work] |
| $r_{exo,growth}$ | 0.68/s | Rate of exocytosis events under growth conditions (20.7 $\mu\text{m}^2/\text{hr}$) [This work] |
| $r_{exo,static}$ | 0.50/s | Static rate of exocytosis events (no net added area) |
| $\sigma_{exo}$ | 0.848 $\mu\text{m}$ | Standard deviation of exocytosis distribution [This work] |
| $\sigma_{endo}$ | 1.838 $\mu\text{m}$ | Standard deviation of endocytosis distribution [This work] |

##### Determination of membrane diffusion and dissociation rates by FRAP

Diffusion coefficients and membrane dissociation constants were determined by fitting FRAP recovery profiles to a 1D model (Fig. S4). The method relies on the different effects of diffusion and association/dissociation on the recovery, with diffusion providing smoothing of profile over time (*65*) and association/dissociation contributing to uniform recovery.

###### Experimental FRAP profiles

Experimental fluorescence intensity profiles were acquired after photobleaching of a rectangular region and imaging, on the top surface of *S. pombe* cells (Fig. S4A; see experimental details in the experimental methods). We corrected for photobleaching after the initial FRAP event by multiplying all intensities by *e^kt^* after background subtraction, where *k* is the exponential decay constant of the average intensity of a neighboring non-bleached cell. From these photobleach-corrected images we determined the intensity profile *I*(*x,t*) by defining a rectangular region of interest along the long axis of a bleached cell with width ~2 μm, excluding the region close to the cell tips, and calculated the average intensity along this line at points spaced 0.0425 μm (1 pixel) apart through the bleaching recovery using ImageJ’s getProfile function (Fig. S4B).

###### Model with diffusion and association/dissociation used for fitting

To match the experimental bleach recovery, we modelled the evolution of the concentration profile of a membrane-associated protein, *c_m_*(*x, t*), along a 1D line with reflecting boundaries at positions *x_a_*, *x_b_*, defining the size of the finite membrane reservoir (approximately the location of cell poles and beyond, to include the back side of the cell). The x-dimension in the model corresponds to the long axis of the cell. We assumed *c_m_*(*x, t*) obeys the following equations, which include both the effects of diffusion and membrane binding/unbinding:

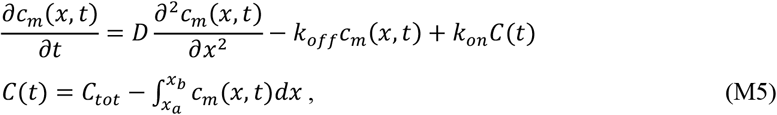

where *c_m_* is concentration on the membrane *k_off_, k_on_* is the number of proteins in the cytoplasm, *C_tot_* is the total number of proteins, and *k_off_*, *k_on_* are the membrane dissociation and association rate constants. We do not write a separate diffusion equation for the cytoplasm, assuming that cytoplasmic diffusion is sufficiently fast. We also specifically consider the limit where most of the proteins are on the membrane: *C*(*t*) ≪ *C_tot_*, (equivalently, the limit of sufficiently large *k_on_*). The Green’s function describing the probability for a single protein being at position *x* at time *t*, given that it was at position *x*_0_ at time *t* = 0 is

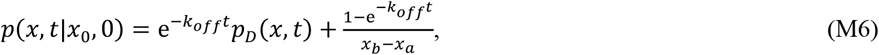

where *p_D_*(*x, t*) accounts for the probability of the particle position due to diffusion alone

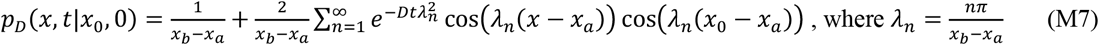

###### General procedure of fitting experimental I(x,t) profiles

Assuming *c_m_*(*x, t*) is proportional to the experimental *I*(*x, t*), we solve for *c_m_*(*x, t*) given an initial condition *c_m_*(*x*, 0) that is proportional to the intensity profile imaged immediately after the photobleaching, *I*(*x*, 0). We then fit the model to both *x* and *t* after photobleaching simultaneously, using SciPy’s optimization curve fitting to get a single best fit value for *D* and *k_off_* as described below. The sum in Eq. (M7) was carried up to *n* = 500. We checked that replacing this by 250 did not have a significant effect on the values determined for *D* and *k_off_* for our experimental data.

###### *Evaluation of initial intensity distribution and simulation boundary positions x_a_* and *x_b_*

The intensity profile is divided into three regions: left, middle, and right (Fig. S4B). The intensity in the middle region, defined to be between *x_d_* and *x_e_* where *x_a_* ≤ *x_d_* ≤ *x_e_* ≤ *x_b_*, is determined by the intensity profile 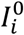 immediately after photobleaching, evaluated at pixels *i* of the projected profile, corresponding to positions *x_i_* (Fig. S4B). The middle region is therefore treated as a sum of *n* = Δ*x*/(*x_e_* – *x_d_*) separate integrals, where Δ*x* is the pixel size. The lengths of the left and right flanking regions are both equal to *l_F_*, with *l_F_* = *x_d_* – *x_a_* = *x_b_* – *x_e_*. We average the intensity of the pre-bleached cell over the middle region and set this as the initial intensity of the two flanking regions, 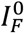. The value of *l_F_* determines the uniform intensity at long times, *I*_∞_, according to mass conservation:

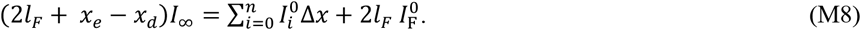

The long time intensity *I*_∞_, or equivalently *l_F_*, is determined via the fitting procedure. In mutants where the *l_F_* value drifted towards 0 or large numbers, it was restricted to be in the range 2-10 μm.

###### Evolution of concentration on a discrete lattice starting from a given initial condition

The equation describing the concentration or intensity as a function of position *x_i_* corresponding to pixel *i*, and time is found by integration, 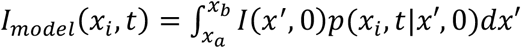, which, using Equations (M6) and (M7) gives:

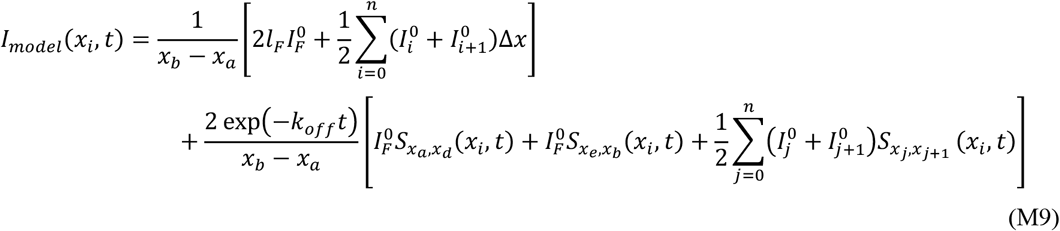

where 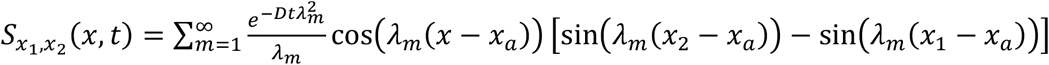 and *λ_n_* is given in Equation (M7). Here, the initial intensity *l*(*x*, 0) at *x_i_* < *x* < *x*_*i*+1_ was assumed to be given by 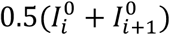. An example of a fit over time to the experimental FRAP data for a cell expressing 1xRitC-GFP using this expression is shown in Fig. S4C.

###### Accuracy of determining D and k_off_

The fit to each FRAP profile determines the single best fit values for *D* and *k_off_* for each cell, corresponding to the largest *R*^2^ value that is a measure of the goodness of the fit. These two values are not independent of each other: since both contribute to recovery, increasing the value of one parameter can be partly compensated by a decrease in the other (even though they influence the evolution of the profile differently, depending on the size of the bleached region). Additionally, one of the two parameters may be too small and thus dominated by the other. These complications are reflected in a large scatter of values for *D* and *k_off_* among cells (Fig. 3A,D and 4D). To check that despite this scatter, averaging over cells is meaningful, we developed a global procedure to determine the best *D* and *k_off_* simultaneously for all cells. For each cell of a given cell type, we varied *D* and *k_off_* over six orders of magnitude and calculated the *R*^2^ value. We then plotted the *R*^2^ value as function of *D* and *k_off_*, averaged over all cells of the same cell type (Fig. S4D-F). The best fitting regions indicated by high values of *R*^2^ typically form a pair of branches: one horizontal (constant *k_off_*) and the other vertical (constant *D*) which intersect to form a curved bend. The median of the best fit value for *D* and *k_off_* as determined by the 1D model is indicated by a red circle. For most photobleached proteins this highest average *R*^2^ (indicated by a blue circle) is close to the median of the best fit values from the 1D model. One case does not show particularly good consistency (1x cc-CBD), where the highest average *R*^2^ is at very low values of *k_off_*), suggesting that the values determined for *D* and *k_off_* for these proteins is not reliable. Vertical or horizontal regions of high *R*^2^ that extend beyond the boundaries of the graphs also indicate that the parameter varied along such a region may only be determined up to an upper limit. They typically correspond to scattered *D* and *k_off_* values in Fig. 3A,D and 4D. Finally, the line defining the ridge of the pair of branches that intersect to form a curved bend in Fig. S4D,E is plotted as continuous line in Fig. 3A,D and 4D to indicate the different combinations of *D* and *k_off_* that provide similar overall recovery.

###### Simulations to validate FRAP fitting procedure

One limitation of our FRAP fit method to extract *D* and *k_off_* is that we neglect the diffusion along the radial cylindrical direction. To examine the implications of this limitation, as well as another check of our method, we simulated FRAP experiments accounting for diffusion and membrane association/dissociation of particles on the surface of a cylinder with reflecting boundary conditions at the ends. We used a similar method as used for the sphere model with 50,000 particles on the surface of a cylinder with radius 2 μm and length 12 μm. As with the discrete sphere model, we fix *k_on_ =* 0.01 *s*^−1^. The displacement is calculated as 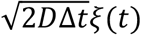 independently in both directions in the plane tangent to the cylinder surface, where Δ*t* is the timestep determined from the Gillespie algorithm and *ξ*(*t*) is a random Gaussian distributed variable with zero mean and unit variance. The displacement along the long axis of the cylinder needs no modification, but the displacement in the radial direction is projected down onto the cylinder surface while maintaining the distance as with the sphere model. To ensure that the radial diffusion is properly implemented, we measure the change in the azimuthal angle, 〈*φ*(*t*)^2^〉, as a function of time up to a time step of 1000 s, similar to Fig. S9A.

The bleaching recovery was measured over a region of width 2.36 μm which is similar to the average experimental width of experimental cells. We took snapshots of particle positions every 1 s for a total of 500 s. To simulate initial photobleaching we label particles on the surface which are within a certain region at a given frame number as being bleached. This simulated bleach region is rectangular on the surface of the cylinder with a long axis of 3.315 μm and an arc length of half the circumference. To calculate the simulated intensity profile, *I_sim_*(*x, t*), we counted the number of particles in this region over time that were not labelled as being bleached in the initial frame and further assume that the intensity is proportional to the number of particles. To get better averages for the bleaching recovery, we simulate box photobleaching at each snapshot and follow the *k* independent recovery trajectories from their respective bleached frame, where *k* is the number of snapshots. We then average the intensity curves for each bleaching recovery at the same amount of time passed since the bleaching frame to obtain 〈*I_sim_*(*x, t*)〉. We then use 〈*I_sim_*(*x, t*)〉 up to a time of 250 s as the input for the 1D model to predict *D* and *k_off_*. When using *D* = 10^−3^ μm^2^/s and *k_off_* = 10^−2^ /s for input parameter values of the discrete simulations, which are approximately in the range of 2xRitC-GFP, there is good agreement between the values inputted into the simulations i.e. the values determined by the *R*^2^ plot (red circle), and the values determined by the 1D model (Fig. S4D). This result indicates that the 1D model can correctly predict *D* and *k_off_* values. For input parameters with zero diffusion (*D* = 0, *k_off_* = 10^−2^ /s) and zero membrane unbinding (*D* = 10^−3^ μm^2^/s, *k_off_* = 0) shown in Fig. S4G the red dashed line indicates the nonzero input parameters are accurately determined, however a small nonzero value is assigned to the other parameter.‘

###### *Comments on measured D* and *k_off_ values*

Even though the estimated values of *D* or *k_off_* can be partly compensated by an inverse change in the other, the FRAP recovery fits to the 1xRitC, 2xRitC, 3xRitC constructs do indicate that the main parameter responsible for slowing dynamics with increasing number of RitC domains is *k_off_* (Fig. S4E). This trend most likely indicates the cooperative nature of the dissociation process for constructs with multiple binding sites (*66*). Our results also point to a potential strong reduction of the diffusion coefficient with increasing number of RitC domains, however more precise measurements are needed to resolve if such a reduction is consistent with a weak size dependence of *D* expected for dilute systems (*67, 68*) or else if this is an indication of crowding mechanisms and microdomain formation in the fluid membrane component (*69, 70*). Interestingly, our measurements also indicate slower diffusion of 3xsfGP-1xRitC compared to sfGP-1xRitC. This observation could indicate that cytoplasmic drag might dominate diffusion of the 3xsfGP construct; further studies could explore if this diffusion reflects size-dependent frictional interaction with cytoplasm adjacent to inner membrane leaflet or else if it is due to oligomerization and related processes.

#### EXPERIMENTAL MATERIALS AND METHODS

##### Media and growth conditions

For microscopy experiments, cells were grown to exponential phase in Edinburgh minimal medium (EMM) supplemented with amino acids as required, unless indicated otherwise. For experiments in Fig 1, Fig 3B-C, Fig S1 and Fig S3, cells were first precultured in 3 ml of EMM in dark conditions at 30 °C for 6–8 h, exponentially growing precultures were diluted (optical density [O.D.]_600nm_ = 0.02) in 10 ml of EMM and incubated in dark conditions overnight at 30 °C, except for *ypt3-i5* and control strains that were cultured at 25 °C in YE. In order to allow proper aeration of the culture, 50 ml Erlenmeyer flasks were used. For experiments in Fig 3D-E, Fig 4, Fig S4 and Fig S6-8, *S. pombe* cells were grown at 25°C. All live-cell imaging was performed on EMM-ALU agarose pads, except for the experiment shown in Fig 1E, for which lectin coated 96-well plates were used.

The optoGAP strain (Fig 4) was generated and grown continuously in light conditions (details on strain generation below). To compare cell morphologies in light and dark conditions (Fig 4E), the strains were grown on YE plates over 2 days at 30°C in the presence of light. Cultures in EMM-ALU were then prepared and split in incubators with or without light and diluted over 4 days to maintain exponential growth until imaged. The strains grown in the dark were not exposed to any light source until the moment of imaging. From these cultures, half of the cell material was used for AiryScan imaging and half was used for Calcofluor imaging. To study the dark-to-light transition (Fig 4F), cultures grown in the dark as above were shifted to light conditions at the microscope during live-cell imaging performed on EMM-ALU 2%-agar pads using blue light exposure at 5 focal planes through the cell depth every 2 minutes. For the drop-test assay (Fig 4J), cultures were grown in the light as above, adjusted to OD_600_ = 0.3, subjected to 5x serial dilutions, and 5μl of each dilution was spotted on YE plates and grown at 30°C under light or dark conditions.

##### Strain construction

Strains used in this study are listed in Table S2 at the end of this section. Standard genetic manipulation methods for *S. pombe* transformation and tetrad dissection were used. Gene tagging was performed at endogenous genomic locus at the 3’ end, yielding C-terminally tagged proteins, as described (*71*). GFP-Ypt3 protein was ectopically expressed in addition to the endogenous gene by transforming a WT strain with PmeI-linearized pBSII(KS^+^)-based single integration vector (pSM2366-3’UTR^*ade6*^-GFP-Ypt3-hphMX-5’UTR^*ade6*^) targeting the *ade6* locus (*72*). Gene tagging, gene deletion, and plasmid integration were confirmed by diagnostic PCR for both sides of the insertion. The construction of plasmids and strains expressing the CRY2-CIB1 optogenetic system was done as described in (*2*).

To generate the RitC strain series, a synthetic gBlock DNA fragment was ordered (Integrated DNA Technologies, Inc.) containing four RitC repeats separated by linker sequences with restriction sites for the generation of 1x, 2x, and 3xRitC constructs. For all ordered synthetic gBlocks with repeat sequences codon usage was modified between repeats to allow successful synthesis. The RitC constructs are N-terminally tagged with sfGFP through sub-cloning in the stable integration vector (S.I.V.) backbone pAV0327 for *ura4* integration under expression from *act1* promoter and with *tdh1* terminator ((*72*); Addgene ID: 133488). Plasmid numbers and descriptions are in Table S3 at the end of this section. Plasmids maps are available upon request. 1xRitC was additionally tagged N-terminally with mCherry in the S.I.V. pAV607 for *ade6* locus integration ((*72*); Addgene ID: 133504). The 3xsfGFP-1xRitC tagging was performed on pAV327 backbone carrying one sfGFP sequence and subcloning of a synthetic gBlock with 2 additional sfGFP sequences and the RitC sequence. Plasmids were integrated as described (*72*). Strains with either mCherry-1xRitC (Fig 3D-E) or CRIB-3xmCherry (Fig S6) along with the respective sfGFP tagged RitC construct were generated through mating and tetrad dissection.

For the structure-function analysis of Rga4, fragments were amplified from the full-length gene originally amplified from cDNA using primers listed in Table S4, listed at the end of this section, and first cloned in pSM621 (pREP41-GFP) for C-terminal tagging with GFP. All tagged fragments were subsequently amplified and introduced in S.I.V. for *ura4* single integration pAV327 in *S. pombe* with *act1* promoter driven expression and *tdh1* terminator (Fig 4B, S7B-C, S7E). The cc-CBD and cc-CBD-Δ2 fragments were additionally also expressed under a *pom1* promoter at the *ura4* locus to allow resolution of single clusters by TIRF imaging (Fig S8B-C). All integrations of AfeI linearized plasmids were performed in a *rga4Δ::kanMX* strain (YSM3826). Synthetic geneBlock DNA fragments were ordered for the generation of the multimerized 4xCBD and 4xcc-CBD-Δ2 constructs and introduced in S.I.V.s by In-Fusion cloning (Takara Bio Inc.) (Fig 4B). The CRY2-mCherry-CBD construct was also generated on a S.I.V. backbone for *ura4* integration and driven by *act1* promoter pAV0328 ((*72*); Addgene ID: 133489) (Fig S8A). Immediately after transformation, this strain was grown and stored only in dark conditions in the absence of any light source until the moment of imaging. Point-mutations in Rga4 membrane binding domains 1 and 2 were generated through several rounds of side-directed mutagenesis with primers listed in Table S4 for both fragments and full length Rga4. The mutagenized fragments were introduced in S.I.V.s and transformed at *ura4* locus as described above. The mutagenesis of the full length Rga4 was performed on a plasmid carrying the open reading frame and 112 base pairs of up- and downstream sequence. The entire sequence was excised (BamHI/SalI) and transformed in *rga4Δ::ura4+* background for integration at the endogenous locus, selected by growth on 5-FOA and verified by absence of growth on EMM-AL medium. The strains were genotyped and verified by sequencing for the presence of mutations. Each mutagenized allele was subsequently tagged with GFP-natMX via standard PCR-based WACH approach (*71*) and additionally introduced in *rga3Δ::hphMX rga6Δ::kanMX* background through the standard *S. pombe* mating protocol and tetrad dissection.

The optoGAP was generated on a S.I.V. for *ade6* locus integration (pAV0356, ((*72*); Addgene ID: 133468)) with *act1* promoter driving the expression of CRY2-mCherry-RitC-rga4(aa665-933). Fragments were assembled by standard cloning. The PmeI linearized plasmid was transformed in a *rga3Δ::hphMX rga4Δ::ura4+ rga6Δ::kanMX* strain (YSM3136; (*28*)) and the strain was continuously grown in the presence of light at 30°C, unless otherwise indicated. We note that attempted transformation of these cells in the dark resulted in absence of colonies, suggesting toxicity as subsequently shown in the serial dilution assay (Fig 4J). The optoGAP strain (*rga3Δ::hphMX rga4Δ::ura4+ rga6Δ::kanMX ade6::pact1-CRY2-mCherry-RitC-rga4(aa665-933)::ade6+*) was then transformed with pAV046 for CRIB-3xGFP-kanMX integration at *leu32* locus in light conditions.

BH-search prediction performed at https://hpcwebapps.cit.nih.gov/bhsearch/ with window size for residue averaging of 15 amino acids and values for amino acids set to standard BH parameters.

**Table S2:**
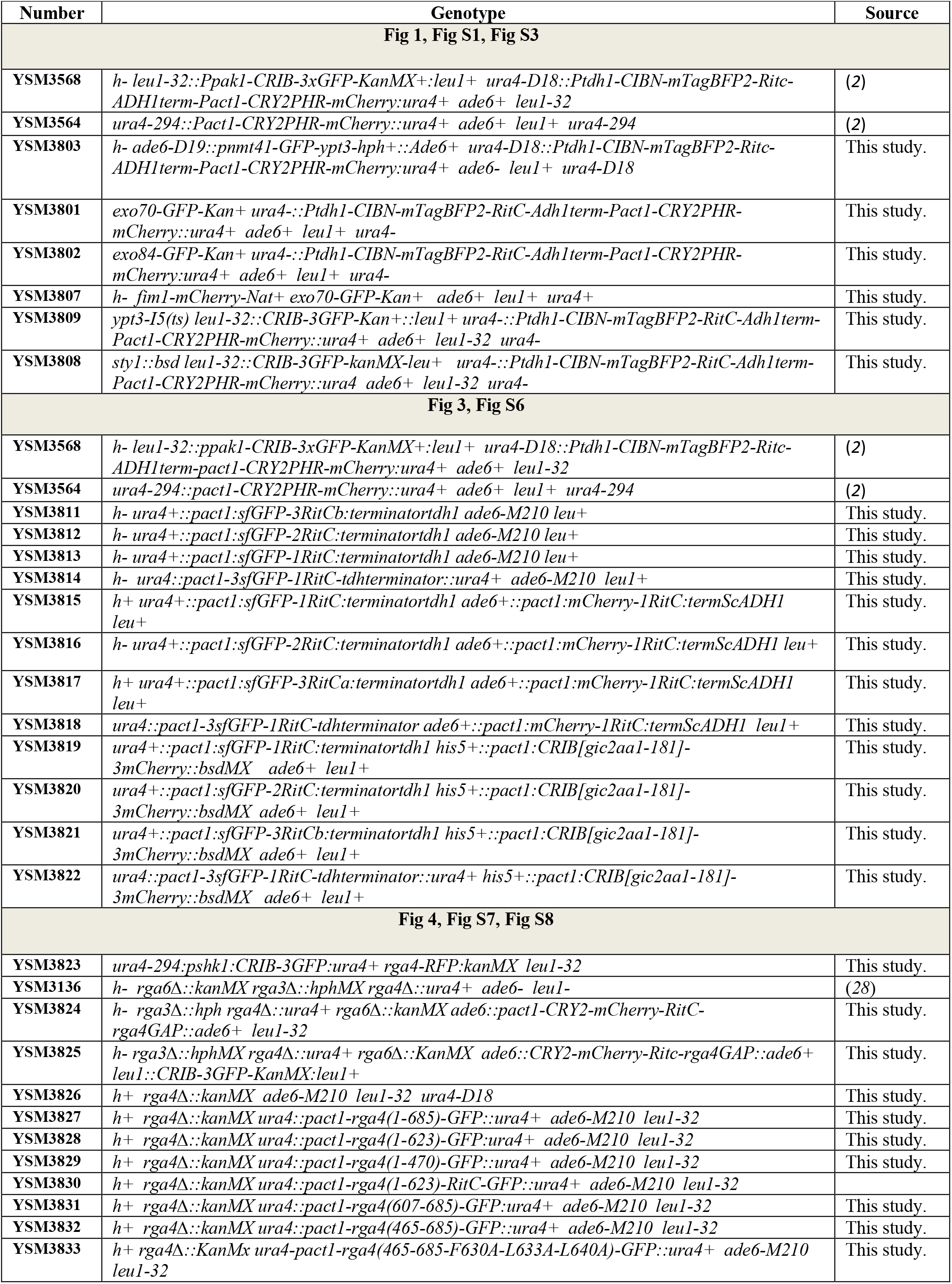
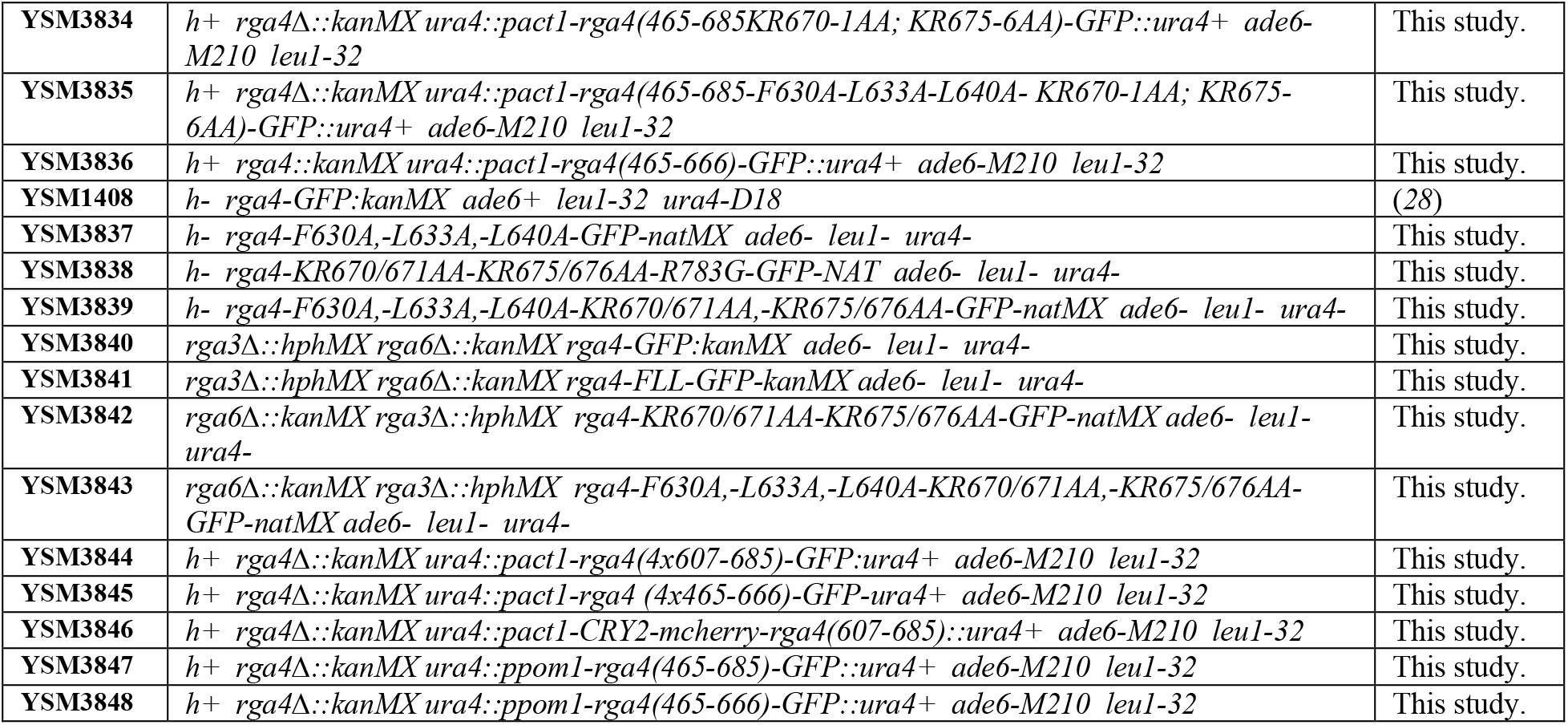
*S. pombe* strains used in this study.

**Table S3:**
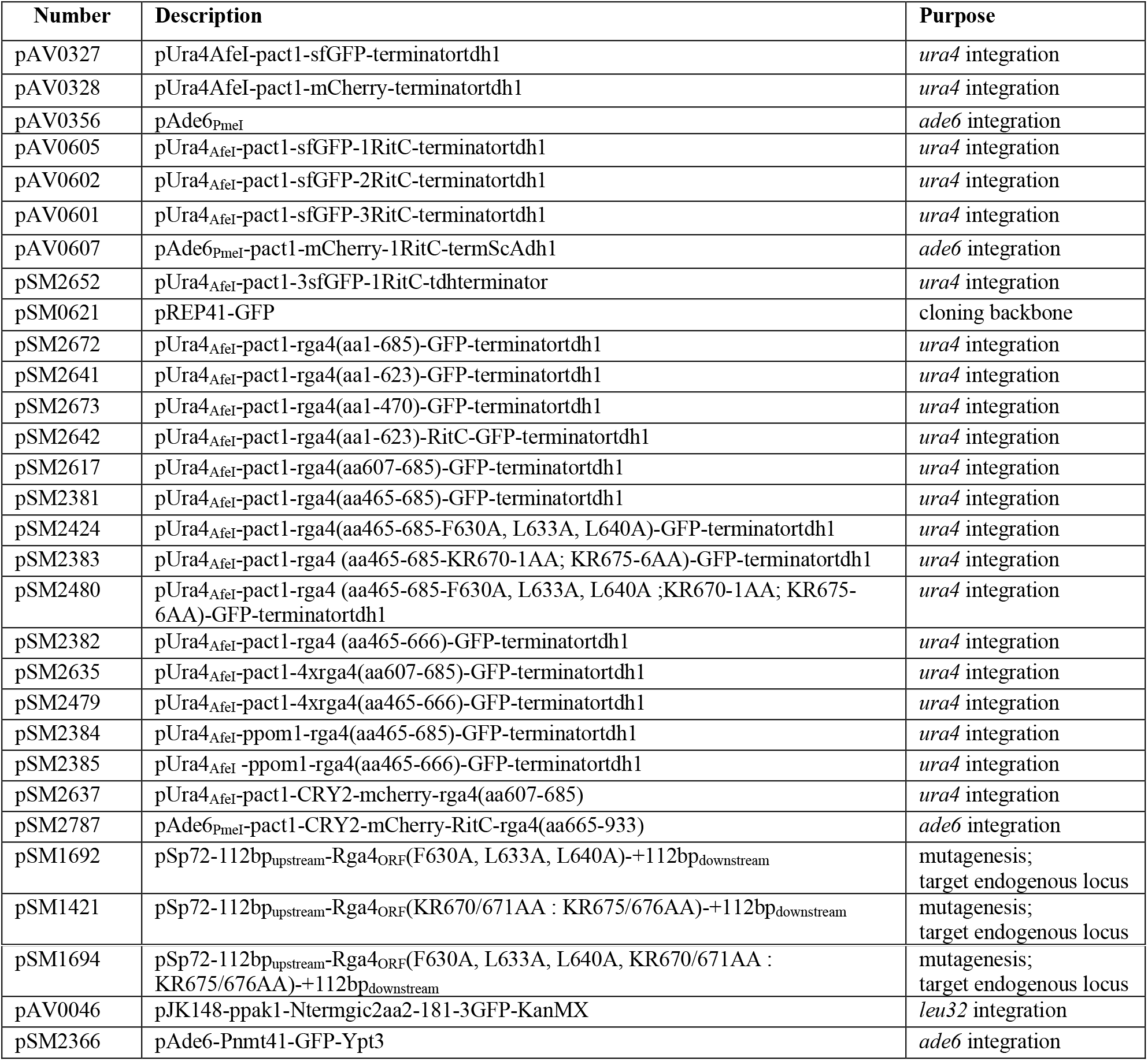
Plasmids used in this study.

**Table S4:**
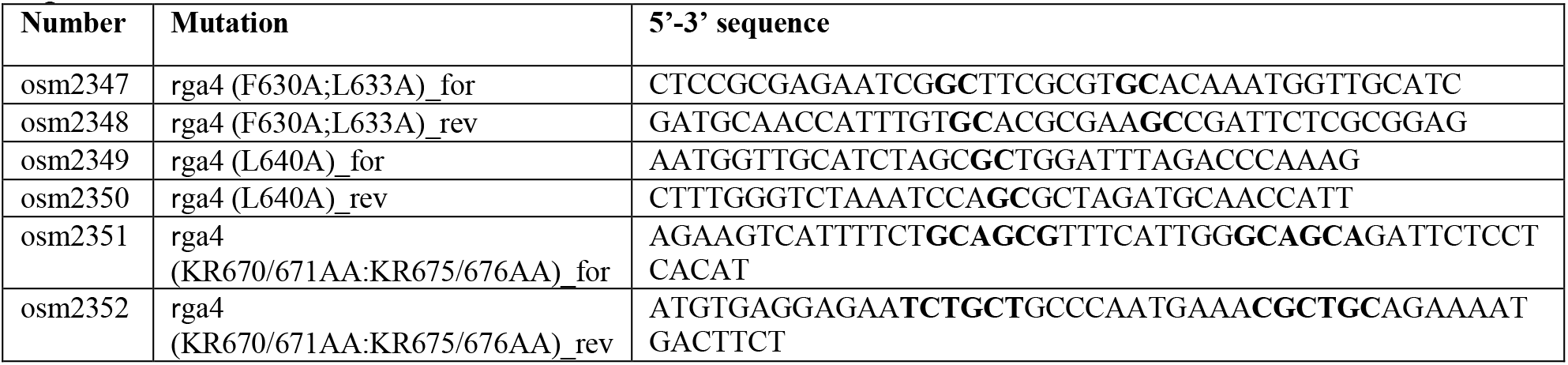
Primers used for mutagenesis. Bold residues indicate changes from the wildtype sequence.

##### Microscopy

###### Spinning-disk microscopy

Images in Fig 1, Fig 3B-C, Fig S1 and Fig S3 were acquired using a spinning disk confocal microscope, essentially as described (*73, 74*). Image acquisition was performed on a Leica DMI6000SD inverted microscope equipped with an HCX PL APO 100X/1.46 numerical aperture oil objective and a PerkinElmer Confocal system. This system uses a Yokogawa CSU22 real-time confocal scanning head, solid-state laser lines, and a cooled 14-bit frame transfer EMCCD C9100-50 camera (Hamamatsu) and is run by Volocity (PerkinElmer). When imaging strains expressing the CRY2-CIB1 optogenetic system, an additional long-pass color filter (550 nm, Thorlabs, Newton, New Jersey, USA) was used for bright-field (BF) image acquisition, in order to avoid precocious photoactivation caused by the white light. In *ypt3-i5* experiments, an objective heater (Bioptechs, Butler, PA, USA) was used for temperature control. Experiments in Fig 1, Fig 3B-C and Fig S1 were carried out using cell mixtures. Cell mixtures were composed of 1 sample strain of interest (the sample optogenetic strain co-expressing the indicated GFP-tagged protein) and up to 2 control strains namely:

1. RFP control: An RFP bleaching correction strain, expressing cytosolic CRY2PHR-mCherry.
2. Drug performance control: A strain expressing specific GFP or RFP markers tagging molecular components targeted by the drug employed in the specific experiment (see Fig S1F and S1I).

Strains were handled in dark conditions throughout. Red LED light was used in the room in order to manipulate strains, to prepare the agarose pads and lectin-coated 96-well plates. Strains were cultured separately. For experiments using agarose pads, exponentially growing cells (O.D._600nm_ = 0.4-0.6) were mixed in 2:1:1 (strain of interest, RFP control, and drug performance control) ratio and harvested by soft centrifugation (2 min at 1,600 rpm). The cell mixture slurry (1 μl) was placed on a 2% EMM-ALU agarose pad, covered with a #1.5-thick coverslip, and sealed with vaseline, lanolin, and paraffin (VALAP). Samples were imaged after 5–10 min of rest in dark conditions. For experiments using lectin-coated 96-well plates, wells were first coated using 100 μl of soy bean lectin (L1395, Sigma-Aldrich) over night. Coated wells were washed 3 times using 200 μl of H_2_O and dried for 4 h at room temperature. Exponentially growing cells (YE liquid media, O.D._600nm_=0.4–0.6) were mixed in 2:1:1 (strain of interest, RFP control, and drug performance control) ratio to a final O.D._600nm_ = 0.1. The cell mixture slurry (200 μl) was placed in lectin-coated wells and left for 45 min in the dark to allow cell-lectin attachment. Finally, cells were washed 3 times with 200 μl YE and 200 μl fresh YE liquid media was added for imaging.

<u>CRY2PHR signal depletion at the cell tips and cell middle</u>. Spinning disk time-lapse images shown in Fig 1A, Fig S1A, Movie 1 and Movie 2 correspond to middle sections focal planes. Images were acquired every 30 s for GFP, RFP and bright-field channels. UV channel was acquired every 1.5 min in order to avoid photo-toxicity caused by UV light. Lasers were set to 100%; shutters were set to sample protection; and in all instances, the GFP channel was imaged first (for CRY2 photo-activation) and then the additional channels. RFP exposure time was set to 200 ms, whereas the GFP exposure was set to 1 s. Cells were monitored in these conditions for 20 min.
<u>CRY2PHR tip depletion dynamic and cell-side accumulation dynamic analyses.</u> Strains co-expressing CRIB, Ypt3, Exo84 or Exo70 GFP-tagged proteins and the CRY2-CIB1 system were monitored in order to assess tip depletion dynamics. In these experiments an RFP control strain was added to the cell mixtures. Results of these experiments are shown in Fig 1B, Fig 3B-C, Fig S1B-E. Lasers were set to 100%; shutters were set to sample protection; and in all instances, the GFP channel was imaged first and then the RFP channel. RFP exposure time was always set to 200 ms, whereas the GFP exposure time varied depending on the monitored protein. Cells were monitored in these conditions for 285 s and images were acquired every 15 s.
<u>Tip depletion dynamics upon disrupting exocytosis using Brefeldin A.</u> For Brefeldin A (BFA; B6542, Sigma-Aldrich) experiments shown in Fig 1C and Fig S1F-G cells co-expressing CRIB-3GFP or GFP-Ypt3 and the CRY2-CIB1 system were imaged. An RFP control strain and a strain expressing Anp1-GFP were mixed in this experiment. Anp1-GFP is a Golgi marker that in normal conditions forms prominent puncta (Fig S1F, left). Upon blocking of endomembrane trafficking due to BFA treatment, Anp1 puncta disappear and the protein re-localizes to the ER (Fig S1F, right). BFA (diluted in ethanol, final concentration 300 μM) was added to the cell mixture slurry. Cells were placed on top of 2% EMM-ALU + BFA 300 μM agarose pads to be imaged. Lasers were set to 100%; shutters were set to sample protection; and in all instances, the GFP channel was imaged first and then the RFP channel. RFP exposure time was always set to 200 ms, whereas the GFP exposure time varied depending on the monitored protein. Cells were monitored in these conditions for 285 s and images were acquired every 15 s.
<u>Tip depletion dynamics upon disrupting exocytosis using *ypt3-i5* ts allele.</u> For *ypt3-i5* experiments shown in Fig 1D and Fig S1H, *ypt3-i5* and *ypt3+* cells co-expressing CRIB-3GFP and the CRY2-CIB1 system were imaged. In these experiments an RFP control strain was added to the cell mixtures. Cells were cultured at 25 °C throughout the experiment and placed on 2% EMM-ALU agarose pad sealed with VALAP. 2 sets of pads were prepared, one to be imaged at 25 °C and another one to be imaged at 36 °C. Prior to imaging, pads were incubated for 30 mins either at 25 °C or 36 °C and imaged at the respective temperatures by setting the objective heater accordingly. Lasers were set to 100%; shutters were set to sample protection; and in all instances, the GFP channel was imaged first and then the RFP channel. RFP exposure time was always set to 200 ms, whereas the GFP exposure was set to 1 s. Cells were monitored in these conditions for 285 s and images were acquired every 15 s.
<u>Tip depletion dynamics upon disrupting endocytosis using Latrunculin A.</u> For Latrunculin A (LatA; BML-T119-0500; Enzo Life Sciences) experiments shown in Fig 1E and Fig S1H-I *sty1*Δ cells co-expressing CRIB-3GFP and the CRY2-CIB1 system were imaged using lectin-coated 96-well plates. An RFP control strain and a strain co-expressing Exo70-GFP and Fim1-mCherry were mixed in this experiment. Fim1-mCherry is an actin patch marker that in normal conditions forms prominent puncta around the cell periphery (Fig S1I, top-left). However, upon actin depolymerization due to LatA treatment, Fim1 puncta disappear and the protein localizes ubiquitously in the cytoplasm (Fig S1I, top-right). LatA (diluted in DMSO, final concentration 50 μM) was added to 200 μl YE liquid media. Lasers were set to 100%; shutters were set to sample protection; and in all instances, the GFP channel was imaged first and then the RFP channel. RFP exposure time was always set to 200 ms, whereas the GFP exposure was set to 1 s. Cells were monitored in these conditions for 285 s and images were acquired every 15 s.
<u>Cortical distribution profiles of exocytosis (Exo70-GFP) and endocytosis (Fim1-mCherry) and counting endocytic events at the cell cortex.</u> A strain expressing endogenous tagged Exo70-GFP and Fim1-mCherry was monitored in order to assess their cortical distribution at the cell tips. Results of these experiments are shown in Fig S3. Lasers were set to 100% and shutters were set to sample protection. Cells were monitored in these conditions for 300 s and images were acquired every 15 s (21 timepoints).

###### Airyscan imaging

All images in Fig 3-4, S4 and S6-8 (except S7D and S8B) were acquired on a Zeiss LSM 980 system with Plan-Apochromat 63x/1.40 Oil DIC objective and acquired by the ZEN Blue software (Zeiss). Imaging was set in super-resolution mode with frame bidirectional maximum speed scanning. Laser power was kept at < 0.4%, with pixel time around 6.6 μs and frame acquisition time <2s per channel. All other settings were optimized as recommended by the ZEN Blue software. In Fig 4A, z-stacks were set at optimal interval with 0.16 μm spacing over a total of 4μm from the medial plane of the cell. All other imaging shows medial focal planes. Frames for movie S5 were acquired with 488nm and 561nm lasers every 5 min and movie 8 and Fig 4F was acquired with 488nm and 561nm lasers every 2 min.

FRAP was performed on cells on a microscope slide (maximum imaging time of 5 min) by finding the focal plane closest to the cover slip and defining a rectangular ROI in the middle of the visible signal (see Fig S4A). 2 snapshots were taken prior to bleaching, which was done using 20% laser power and a 5 time repeat of the bleach zone. To determine the depth of the bleached signal immediately post-bleaching, z-stacks optimally spaced at 0.16μm interval were acquired across the entire depth of the cell, which allowed to calculate that bleaching occurs over a depth of 2.0μm-2.5μm. For 2xRitC and 3xRitC, recovery was imaged for 5 min, every 10s; 1xRitC and Rga4 fragments were imaged at maximum speed every 729ms for 1 min. For FRAP of CRY2-CIBN in the light-activated state, cells were first found in focus using the 561nm laser after which they were illuminated with transmitted white light for 45s and a snapshot of CRY2 to CIBN recruitment at the mid-plane of the cell was taken. The FRAP was then performed as described above at the focal plane closest to the cover slip over a 5 min recovery time, acquired every 10s, at each time point acquisition was performed first with 561nm laser, followed by 488nm laser for a total frame acquisition time of 900ms. The recovery for the FRAP of CIBN-GFP alone was recorded over 1 min, every 700ms maximum speed imaging. All AiryScan images were processed in Zeiss Zen Blue software.

###### Other imaging

For cell length measurements (Fig 4I), cells were stained with 2μg/ml f.c. calcofluor (Fluorescent brightener 28, F3543, Sigma) and imaged with a Leica epifluorescence microscope (63X magnification). TIRF microscopy (Fig S8B-C) was performed on a DeltaVision OMX SR imaging system, equipped with a 60x 1.49 NA TIRF oil objective (oil 1.514), an illumination pathway for ring-TIRF and a front illuminated sCMOS camera size 2560×2160 pxl (manufacturer PCO). Imaging settings were: 512×512 pxl field of view, 100ms exposure time, laser power of 20% with TIRF angle 488nm at 86.7°. Samples were placed on a 0.17 +/- 0.01 mm thick glass slide and imaged within 15 minutes. Imaging was performed every second over a 60s imaging period.

##### Image analysis of concentration profiles

All image-processing analyses were performed with Image J software (http://rsb.info.nih.gov/ij/). Image and time-lapse recordings were imported to the software using the Bio-Formats plugin (http://loci.wisc.edu/software/bio-formats). Time-lapse recordings were aligned using the StackReg plugin (http://bigwww.epfl.ch/thevenaz/stackreg/) using the rigid body method. All optogenetic data analyses were performed using MATLAB (R2019a), with scripts developed inhouse. Figures were assembled with Adobe Photoshop CC2019 and Adobe Illustrator 2020.

###### Correlating CRY2PHR-mCherry tip depletion dynamics with endogenous GFP-tagged protein signal intensities at cell tips

CRY2PHR-mCherry tip depletion dynamics was monitored in strains co-expressing CRIB (N = 3, n = 275 cells), Ypt3 (N = 3, n = 184 cells), Exo84 (N = 3, n = 290 cells) and Exo70 (N = 3, n = 281 cells) GFP-tagged proteins (sample strains) in normal conditions (Fig 1B and S1C-E); in strains co-expressing CRIB-3GFP and GFP-Ypt3 treated with BFA (CRIB: N = 3, n = 219 cells; Ypt3: N = 3, n = 230 cells) or ethanol as control (CRIB: N = 3, n = 127 cells; Ypt3: N = 3, n = 136 cells); in *ypt3-i5* CRIB-3GFP and *ypt3+* CRIB-3GFP strains at 25 ° C (*ypt3+* CRIB-3GFP: N = 3, n = 191 cells; *ypt3-i5* CRIB-3GFP: N = 3, n = 132 cells) and 36 °C (*ypt3+* CRIB-3GFP: N = 3, n = 229 cells; *ypt3-i5* CRIB-3GFP: N = 3, n = 204 cells); and in *sty1*Δ CRIB-3GFP strain treated with LatA (N = 3, n = 276 cells) or left untreated (N = 3, n = 293 cells). Tip depletion was assessed upon photoactivation of the CRY2-CIB1 system by recording the RFP fluorescence intensity over an ROI that was 12 pixels wide by 25 pixels long (≈ 1 μm by 2.075 μm) and drawn perpendicular to the cell tip cortex over 20 timepoints, as shown by the blue arrow in the scheme on the right. The RFP and GFP fluoresce profiles across the plasma membrane were recorded over time. The GFP signal was used to correlate the presence of endogenous proteins with the CRY2PHR-mCherry tip depletion dynamics. Note that in these sets of experiments the initial timepoint was not included in the analysis, since CRY2PHR does not yet achieve its maximal plasma membrane recruitment (see Fig 3B). Thus, the second time point was set as the initial timepoint (t_0_) in these experiments (19 timepoints in total). This corresponds to t = 0 s in Fig 3B.

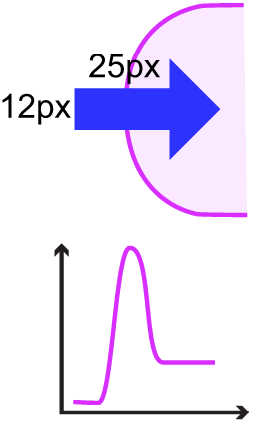

To derive photobleaching correction coefficients, the average camera background signals (*Bckg*) from 6 cell-free regions were measured; and fluorophore bleaching was measured from entire cells, using RFP controls for RFP and sample cells for GFP.

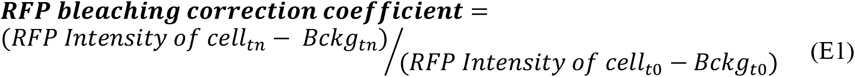

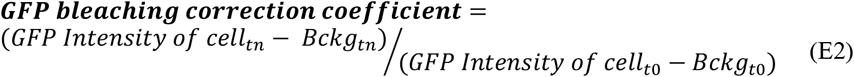

where *RFP Intensity* and *GFP Intensity* of cell stands for the signal measured from entire RFP control and sample cells, respectively; *Bckg* stands for the average fluorescence intensity of 6 independent cell-free regions; t_n_ represents a given time point along the time course of the experiment; and t_0_ represents the initial time point (*n* = 19 time points). These coefficients were corrected by a moving average smoothing method (moving averaged values = 5). GFP and RFP profiles of sample cells were independently analysed as follows. First, GFP and RFP signals were background and bleaching corrected, using Eq E1 and E2:

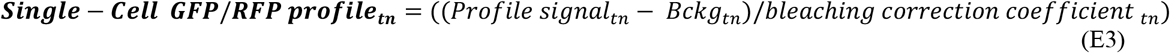

where *Profile signal* intensity represents the GFP or RFP raw profile values across the plasma membrane, *Bckg* stands for the average fluorescence intensity of 6 independent cell-free regions, and t_n_ represents a given time point along the time course of the experiment.

Corrected single-cell GFP and RFP profiles were then aligned to their peaks in order to correct for cell growth and/or cell movement. The RFP peak at t_0_ was used as a reference point to align singlecell RFP and GFP profiles over time. The profiles resulting from the peak alignment were used to calculate the cortical GFP and RFP fluorescence values over time:

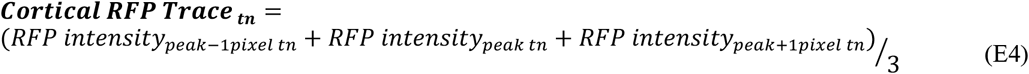

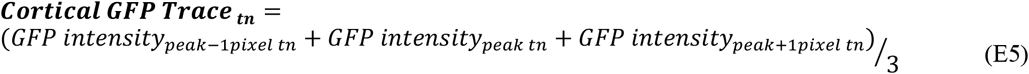

The GFP and RFP fluorescence intensities at RFP_peak pixel position at t0_ ± 1 pixel were averaged in order to extract CRY2PHR-mCherry tip depletion (RFP channel) and GFP-tagged traces across the duration of the time-lapse. From here on, GFP traces were saved and further analyses were performed only for RFP traces. First, the amplitude of the RFP tip depletion was calculated as follows:

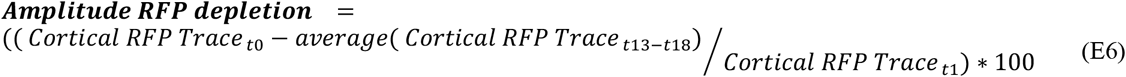

where *Cortical RFP trace t_0_* represents the initial RFP signal at the plasma membrane and *average (Cortical RFP trace t_13-18_*) represents the average signal at the plasma of the last 5 timepoints of the trace. In addition, single-cell RFP traces derived from Eq E4 were also normalized to their own maximum RFP intensity:

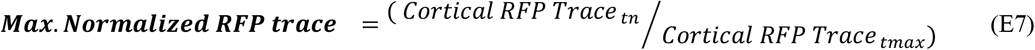

where *Cortical RFP trace t_n_* represents the RFP value of the trace at the timepoint *n* and *Cortical RFP trace t_max_* represent the maximum RFP value of the single-cell trace.

From here on, normalized RFP traces from Eq E7 were sorted according to the amplitude of depletion derived from Eq E6. When the *Amplitude RFP depletion* was lower than 20 %, then the normalized RFP trace from Eq E7 was labelled as “Non-depleted”. These traces are represented in red-coloured dots in Fig 1B and similar cluster plots. Cumulative GFP values of “Non-depleted” RFP traces were calculated as follows:

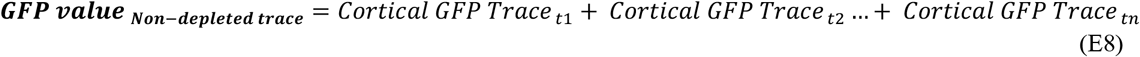

Where *Cortical GFP trace t_n_* represents the GFP value at timepoint n from the GFP trace calculated in Eq E5. Thus, GFP value for “Non-depleted” RFP traces correspond to the sum of GFP values along the 19 timepoints.

In contrast, when the *Amplitude RFP depletion* was greater than 20 %, then the normalized RFP trace from Eq E7 were labelled as “Depleted”. These “Depleted” traces were used to calculate the tip depletion half-times:

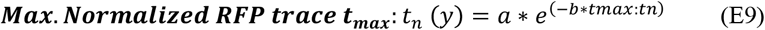

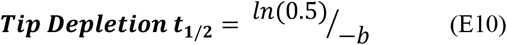

where *Max. NormalizedRFP trace t_max_:t_n_* represent the normalized RFP trace from Eq E7 starting from the timepoint of its maximal RFP value (*t_max_*) to the last timepoint of the trace (*t_n_*) and *t_n_* represents the last timepoint of the trace. If t_max_ corresponds to t_0_, then it means that the tip was depleting from the very beginning of the recording. In contrast, if tmax does not corresponds to t_0_, then it means that the tip depletion started later on. Therefore, a *Depletion delay* variable was also calculated.

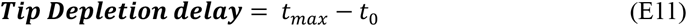

where *t_max_* represents the timepoint at which the maximal RFP value was detected along the single-cell trace and *t_0_* represents the first timepoint of the trace. 67% of traces started depleting from t_0_, 19% from t_1_ and a few % thereafter.

Finally, cumulative GFP values for “Depleted” RFP traces were calculated as follows:

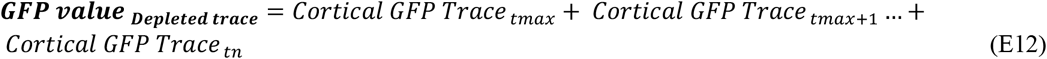

where *Cortical trace t_max_* represents the GFP value of the GFP trace calculated in Eq E5 at timepoint *tmax*. Thus, GFP value for “Depleted” RFP traces correspond to the sum of GFP values from *t_max_* to t_n_ (*n = 19 timepoints*).

After these data analyses, RFP traces were further sorted based on the R^2^ value of the fitting from Eq E9 and the *Tip depletion delay* value from Eq E11. “Depleted” RFP traces matching any of these criteria were discarded from the regression analysis:

1. The depletion delay from Eq E11 is more than 90 s (starts depleting after t7).
2. R^2^ of curve fitting from Eq E9 is lower than 0.7, except in *ypt3-i5* and LatA experiments, where R^2^ > 0.5 was accepted.

Additionally, “non-depleted” RFP traces, which depleted less than 20% but initiated depletion after t14 (delay from Eq E11 is more than 200 s) were discarded, as these may represent tips that may in fact eventually deplete, after the time of the time-lapse.

Finally, the cumulative GFP values from Eq E8 and Eq E12 and the amplitude of RFP depletion from Eq E6 of non-depleted and depleted traces were used to perform a linear regression analysis. The linear regression was performed using the correlation matrix scatterplot function for MatLab developed by John Chow* (*75*).

###### CRY2PHR-mCherry lateral peak formation

The lateral accumulation of CRY2PHR-mCherry at the edges of the depletion zone in a strain coexpressing CRIB-3GFP and the CRY2-CIB1 system (N = 3, n = 268 traces in 134 cells) was monitored by recording the RFP and GFP fluorescence profiles over an ROI that was 3-pixels wide by more than 171-pixels long (≈ 0.25 μm by 14.2 μm) drawn along the cortex of sample cells, as shown by the blue arrow on the scheme on the right. The ROI was drawn from one cell side towards the other cell side, passing across the tip of the cell. The RFP and GFP fluoresce profiles were recorded over time from sample strains. The GFP signal was used to align single-cell RFP and GFP profiles. Because CRY2PHR only achieves its maximal plasma membrane recruitment at the second image acquisition, this was set as t = 0 s. (see Fig 3B).

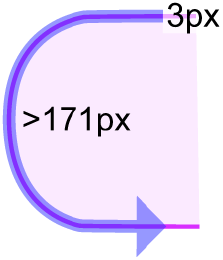

Photobleaching correction coefficients were derived as in Eq E1-E2 and single-cell RFP and GFP cortical profiles were corrected using Eq E3. Single-cell cortical GFP profiles from all the timepoints (*n = 20 timepoints*) were averaged in order to generate an *average Cortical GFP profile*. The peak of the *average Cortical GFP profile* was used as a reference for the center of the cell tip. The peak pixel position of the *average Cortical GFP profile* was fitted to a Gaussian distribution, where the parameter *b* corresponds to the peak position:

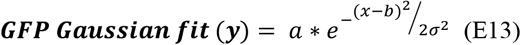

Finally, all single-cell GFP and RFP profiles were aligned to the center of the cell tip, divided symmetrically in two traces and averaged per timepoint (Fig 3B, bottom).

Tip depletion and cell-side accumulation traces were calculated from *single-cell RFP profiles* by averaging the RFP intensities at pixel positions shown in Fig 3B (bottom) for every timepoint. For each cell the average fluorescence value at the cell tip (sampled over 15-pixel length = 1.25 μm), showing depletion over time, and at the edge of the depletion zone (sampled over 10-pixel length = 0.83 μm), showing accumulation over time, was calculated. Average traces are shown in Fig 3B (top).

###### Cortical distribution profiles of exocytosis (Exo70-GFP) and endocytosis (Fim1-mCherry) at the cell tips and cell middle

To experimentally address the relative distribution of endo- and exocytosis, we compared the distribution of the exocyst subunit Exo70 and the actin-binding Fimbrin Fim1, as markers of exo- and endocytic events, respectively (*64, 76–78*). Both markers are enriched at cell poles and division sites, and we examined their distribution at both locations (Fig S3). The distribution profiles were generated from the sum projection images of entire time-lapses (n = 21 timepoints). Exo70-GFP and Fim1-mCherry fluorescence profiles were recorded over ROIs drawn from one cortical cell side towards the other cell side (N = 3, n = 149 profiles) and at the cell middle cortex (N = 3, n = 70 profiles) (see Fig S3A and S3B). ROIs were 6-pixel wide by more than 171-pixel long (≈ 0.5 μm by 14.2 μm) for cell tips and 6-pixel wide by 92-pixel long (≈ 0.5 μm by 7.64 μm) for the cell middle.

Single cell profiles were first corrected for the background signal using Eq E14 and E15.

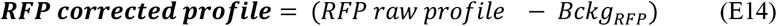

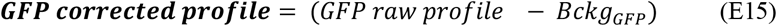

where *RFP/GFP raw profile* stands for Exo70-GFP and Fim1-mCherry profiles generated from ROIs drawn at cell cortex; and *Bckg_RFP/GFP_* stand for the average fluorescence intensity of 6 independent cytosolic regions. *Bckg_RFP/GFP_* serve to set fluorescence threshold present in the cytosol of Exo70-GFP and Fim1-mCherry cells. Due to the ubiquitous Fim1-mCherry cortical localization and GFP auto-fluorescence, *Bckg* ROIs were placed in the interception between the nucleus and the cytosol in the centre of cells co-expressing both markers. Upon background correction, RFP and GFP profile values below the threshold (i.e. < 0 A.U.) were set to 0 A.U. in order to prevent numerical aberrations during the analysis.

Single-cell *corrected GFP/RFP profiles* were then normalized to their maximum and minimum values and fitted to a Gaussian distribution using Eq E13. Finally, single-cell *corrected GFP/RFP profiles* were aligned to their geometrical center, averaged and normalized to the maximum and 0 values (Fig S3).

We note that Exo70-GFP exhibits a narrower distribution than Fim1 -mCherry at growing cell poles (Fig S3A). We observed similar, even more extensive distribution differences in the middle of pre-divisional cells (Fig S3B). These observations show that endocytosis occurs in a region centred over, but wider than, the zone of polarized exocytosis. These findings are in line with previous observations in several fungi (*79–82*). The measured distributions at cell poles were used in the simulations.

###### Estimating endocytic and exocytic rates

To estimate the rate of endocytosis, we counted the number of Fim1-mCherry patches at the cell side of cells co-expressing Exo70-GFP and Fim1-mCherry markers across the duration of the whole time-lapse. We also measured the average fluorescence intensity over the same cell-side regions in sum intensity projection images. These values were used to calculate an average Fim1-mCherry patch fluorescence. We performed this analysis at cell sides rather than cell tips to avoid overlap of endocytic signal at cell tips due to high density, which would lead to an underestimation of the number of events. By dividing the Fim1-mCherry distribution profile described above by the average patch fluorescence value, the average endocytic rate per pixel position across the Fim1-mCherry distribution profile was calculated as follows:

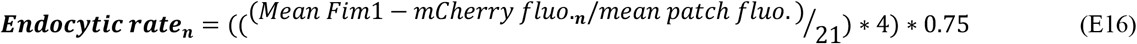

where *Mean Fim1-mCherry fluo._n_* stands for the average RFP fluorescence value at pixel position n; *mean patch fluo*. stands for the average Fim1-mCherry patch fluorescence. The value 21 represents the number of time points that contribute to the average Fim1-mCherry distribution profile. This division aims to estimate the number of endocytic event per image. This is multiplied by 4 to obtain the number of events per minute (imaging interval = 15s). Finally, the result of this operation was corrected by 0.75 in order to correct for the slight over sampling in our images, as the average lifetime of a Fim1-mCherry patch is about 20 s (*78*).

The rate of exocytosis events under growth conditions was calculated from the addition of the above rates of endocytosis and rates membrane addition contributing to cell growth. Cell growth was measured in time-lapse imaging as the difference in length over time between the invariant birth scar location and the pole of the cell, and converted to amount of required membrane to accommodate this growth (see model methods for further details).

The obtained values were used in the simulations and are presented in table S1 above.

###### Cortical distribution profiles of GFP-1RitC, GFP-2RitC, GFP-3RitC and 3GFP-1RitC

Strains co-expressing CRIB-3mCherry and any of the GFP-1RitC (N = 3, n = 266 profiles), GFP-2RitC (N = 3, n = 194 profiles), GFP-3RitC (N = 3, n = 246 profiles) or 3GFP-1RitC (N = 3, n = 274 profiles) constructs were imaged. Cortical distribution profiles were generated from single timepoint middle section images. The RFP and GFP fluorescence profiles were recorded over an ROI that was 3-pixels wide by more than 171-pixels long (≈ 0.25 μm by 14.2 μm) drawn along the cortex of sample cells. The ROI was drawn from one cell side towards the other cell side, passing across the tip of the cell. To correct single-cell profiles for the average camera background signals (*Bckg*), the average signal from 6 cell-free regions was subtracted using Eq E14 and E15. Single-cell *corrected GFP/RFP profiles* were then aligned to their geometrical center and divided symmetrically in two traces starting from the cell pole. To normalize for the differential plasma membrane binding affinity of the GFP-1RitC, GFP-2RitC, GFP-3RitC or 3GFP-1RitC constructs, and compare fluorescence distribution, the area under the curve (AUC) of single-cell *corrected GFP traces* were normalized to 1:

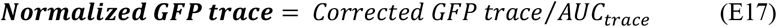

where *Corrected GFP trace* represents the fluorescence profile from the cell pole after background correction and *AUC_trace_* stands for its area under the curve. Finally, *normalized GFP traces* were averaged (Fig 3E).

For the quantification of tip to side fluorescence intensities of the RitC series and correlation to CRIB-3xmCherry (Fig S6), 7 px-wide lines were drawn, one 12 px-long at the tip and two covering the cell sides to measure an average fluorescence signal at cell tip and at cell sides, respectively. Background corrections from the average value of 6 ROIs outside any cell from each individual field of view were performed for each channel. Tip/side ratios were calculated and plotted against CRIB intensity as scatter plots, with R^2^ value calculated from the trend line generated in Excel. Data was collected from three replicate experiments for a total of 100-180 cells per strain.

###### Distribution profile of Rga4 and Rga4 fragments

For the corset profiles of Rga4 in Fig 4A, an 80 px-wide line was drawn across the cell length starting from the cell tip with higher CRIB intensity on a sum projection image obtained from z-stack AiryScan acquisition. Background correction from the average value of 6 ROIs in regions outside cells from each individual field of view was performed.

Cortical profiles for Rga4 fragments (Fig 4C) were manually drawn as 7px-wide cortical lines from cell tip to cell tip along each side of the cell and data was collected using the ImageJ multiplot function. The middle of each profile was identified, the profiles were split in half and aligned to cell tips using MatLab. Background correction from the average value of 6 ROIs outside of cells from each individual field of view was performed. The plots represent data from three replicate experiments with a total of 150-180 cells per strain, shaded area is standard deviation between cells.

TIRF movies were analysed using the TrackMate Fiji plug-in (*83*) and tracking was performed on individual cells. The particle diameter was set to 0.3 μm with maximum gap linking of two frames, and linking range of 0.15 μm. Quality control for spot identification was set at cut-off of 19 and was visually confirmed for each movie.

All boxplots and violin plots were generated using the BoxPlotR web-tool (http://shiny.chemgrid.org/boxplotr/) with definition of whisker extend by Tukey.

###### Analysis of optoGAP

The quantification of optoGAP in Fig 4G shows the number of cells with a clear CRIB zone divided by the total number of cells in both light and dark condition. Quantifications were performed from 3 replicate experiments with a total number of counted cells indicated in the figure. Error bars represent standard deviation between experiments. The CRIB fluorescence intensity profile in Fig 4H was quantified by drawing, for each cell with a CRIB zone, two 7 px-wide profiles along the cell cortex starting at the middle of the cell tip. Data was collected using the multi-plot function of ImageJ and corrected for background from ROIs outside of cells for each individual field of view. For the *rga3Δ rga4Δ rga6Δ* cells without transgene, only cells with a single CRIB zone were considered. Data was collected from three replicate experiments for a total of 150 cells per strain. Shaded area represents standard deviation between cells. The aspect ratio in Fig 4I was calculated from the measured cell length to cell width of dividing cells in three replicate calcofluor experiments for 500-600 cells per strain. Statistics were performed on the averages of the three experiments using a student t-test with equal variances in Microsoft Excel.

###### Cortical kymographs

Kymographs shown in Fig 3C, Fig 4F and Fig S3 were generated with the MultipleKymograph (https://www.embl.de/eamnet/html/body_kymograph.html) ImageJ plugin. In Fig 3C and Fig 4F, a single-pixel-wide ROI was drawn along the cell cortex. In Fig S3, a 6-pixel-wide ROI was drawn along the cell cortex. Fluorescence was averaged to 1-pixel-wide lines to construct the kymographs (parameter Linewidth = 1).

**Figure S1.**
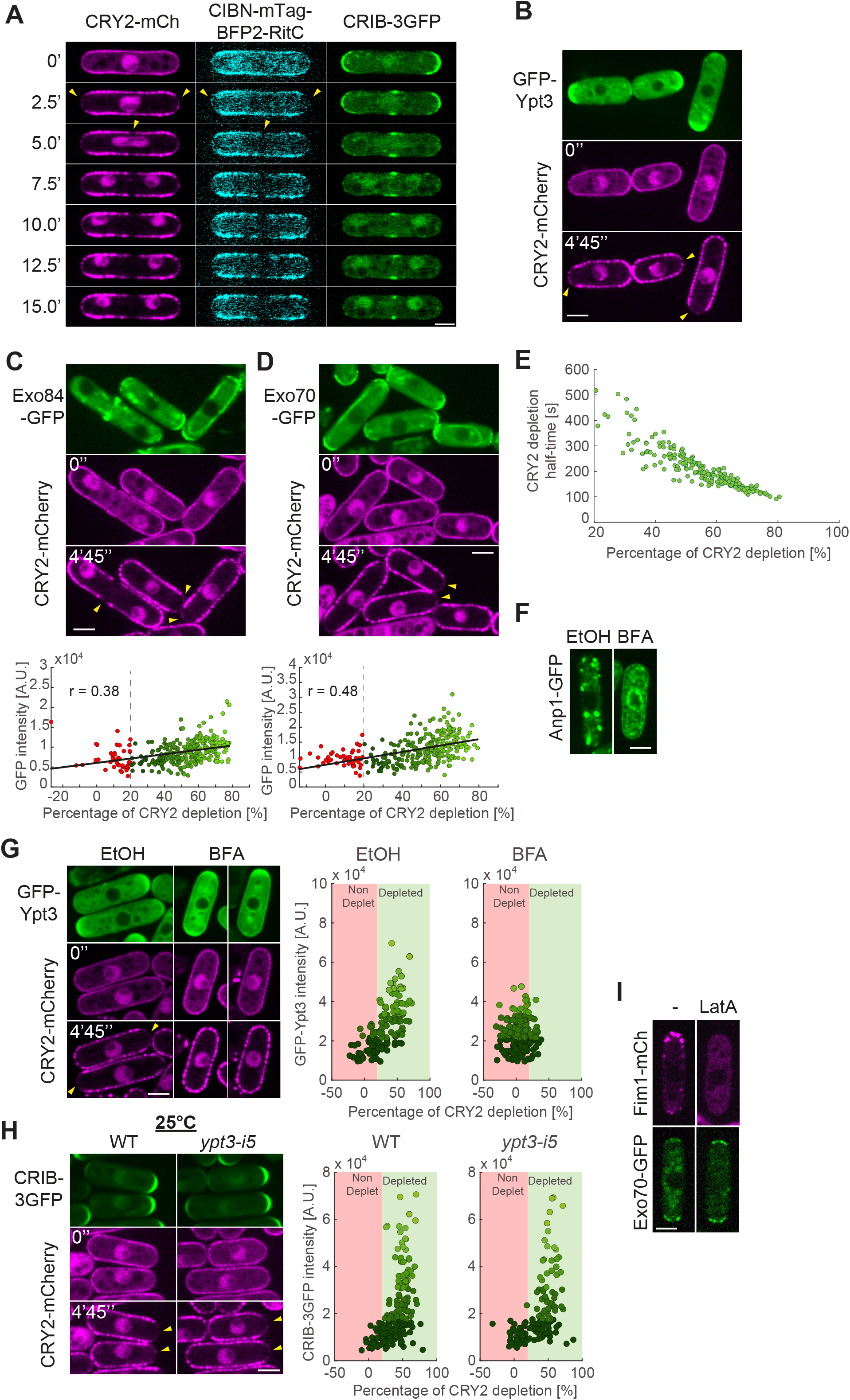
Controls and further evidence of depletion of membrane-associated CRY2 around sites of secretion. **A.** Time-lapse of CRY2-mCherry, CIBN-mTag-BFP2-RitC and CRIB-3GFP in a pre-divisional cell grown in the dark. Time 0 is the first timepoint after illumination. **B.** Time-lapse of CRY2-mCherry and CIBN-mTag-BFP2-RitC redistribution in cells co-expressing GFP-Ypt3 (see Fig 1B for quantification). **C-D.** Time-lapse of CRY2-mCherry and CIBN-mTag-BFP2-RitC redistribution in cells co-expressing Exo84-GFP (C) or Exo70-GFP (D). The bottom graph shows correlation between extent of CRY2 depletion after 5 min and pole GFP intensity. In (B-D), cells were grown in the dark. Time 0 is the first timepoint after illumination. The GFP channel is shown as sum projection over the 5 min time-lapse. The mCherry channel shows individual time points. **E.** Correlation between CRY2 depletion halftime and extent of depletion after 5 min. **F.** Example of Anp1-GFP-expressing cells mixed with cells shown in Fig 1C and panel (G), showing the effect of brefeldin A (BFA) on collapse of Golgi to the ER. **G.** Cells as in (B) treated with 50nM brefeldin A (BFA) or solvent (EtOH). **H.** Cells as in Fig 1D, but at 25°C. In (G-H), the GFP channel is shown as sum projection over the 5 min time-lapse. The mCherry channel shows individual time points. Graphs on the right are as in Fig1C-E with depletion cut-off shown in red and green backgrounds. Data points are coloured with different shades of green according to GFP fluorescence levels. **I.** Example of cell expressing both Fim1-mCherry and Exo70-GFP mixed with cells shown in Fig 1E, showing the effect of Latrunculin A (LatA) on loss of actin patches (but not exocyst localization). Throughout, yellow arrowheads indicate depletion zones. Scale bars are 3μm.

**Figure S2.**
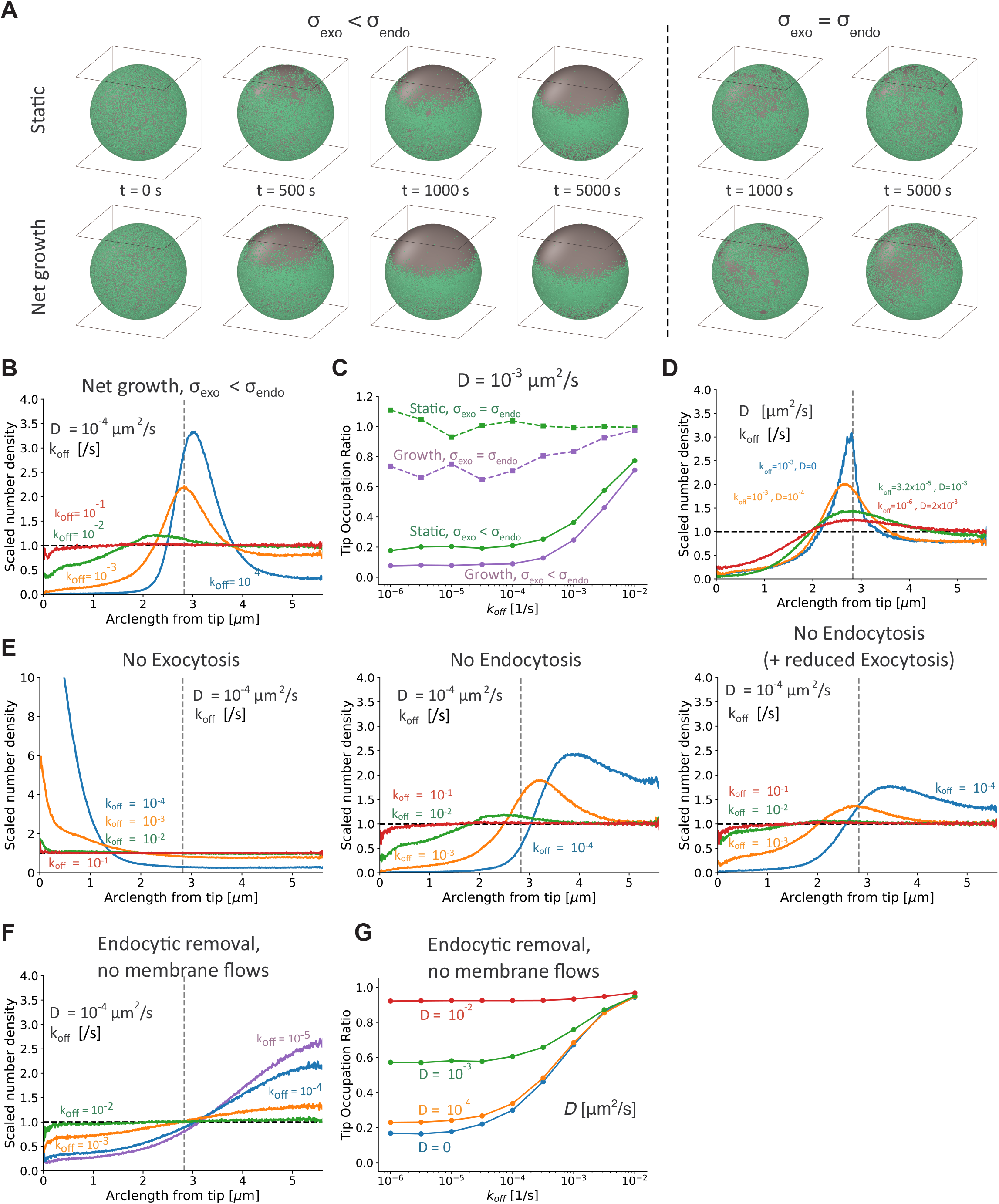
Simulation results and conditions for tip depletion and lateral peak. **A**. Time evolution of particles (green) on the surface of the sphere with *k_off_* = 10^−5^/*s* and *D* = 10^−4^ *μm*^2^/*s* under static conditions (top row; net area added is zero with 0.5 endocytosis events/s and 2.44 endocytosis events/s) and under net growth conditions (bottom row; net area added 20.7 *μm*^2^/*hr* with 0.68 exocytosis events/s and 2.44 endocytosis events/s). The spherical shell representing the cell remains at a constant size; exocytosis and endocytosis only affect the positions of particles on the surface. Left: both rows have *σ_exo_* = 0.85 *μm* < *σ_endo_* = 1.84 *μm* derived from experimental measurements; depletion near the top tip is evident in both cases but occurs faster under growth conditions. Right: both rows have *σ_exo_* = *σ_endo_* = 1.84 *μm*; slight depletion near the tip is only evident under growth conditions. **B.** Scaled number density as a function of arclength for the same simulations as Fig 2E but under growth conditions. Compared to Fig 2E, the lateral peak is shifted further towards the back of the sphere and tip depletion is stronger. **C.** Tip occupation ratio as a function of *k_off_* for static and growth conditions for *σ_exo_* = *σ_endo_* and *σ_endo_* < *σ_end0_* as in panel A, for *D* = 10^−3^ *μm*^2^/*s*. Switching from *σ_exo_* = *σ_end0_* to *σ_ex0_* < *σ_end0_* or from static to growth conditions both result in increased tip depletion. **D.** Increasing diffusion coefficient while keeping the tip occupancy ratio nearly constant reduces the lateral peak height (net growth, *σ_ex0_* < *σ_end0_* as in A). **E.** Scaled number density as a function of arclength, for simulations where exocytosis is turned off/endocytosis remains (left); endocytosis is turned off/exocytosis remains the same as under growth conditions (middle); and endocytosis is turned off/exocytosis is reduced so that the net area addition rate is 20.7 *μm*^2^/*hr* (right). Reference parameters are for net growth, *σ_ex0_* < *σ_end0_* as in (A). Stopping exocytosis leads to tip enhancement for low enough *k_off_*. As these simulations do not account for anticipated slowing down of endocytosis due to reduction of plasma membrane lipid reservoir, we consider them consistent with the results of experiments in Fig 1C-D. Stopping endocytosis maintains tip depletion, in line with experiments of Fig 1E. **F.** Scaled number density as a function of arclength at *D* = 10^−4^ *μm*^2^/*s* under no membrane flow conditions where exocytosis/endocytosis does not move the particles on the sphere surface but endocytosis pulls particles into the internal pool if they are within *R_end0_* of the endocytosis event (using 2.44 endocytosis events/s and *σ_end0_* = 1.84 *μm* as in (A)). Depletion is still evident but the lateral peak near the equator of the sphere disappears. Compare to Fig 2E. **G.** Tip occupation ratio versus *k_off_* under no membrane flow conditions as in (F). Tip depletion is less pronounced compared to simulations with membrane flow in Fig 2D, for the same conditions.

**Figure S3.**
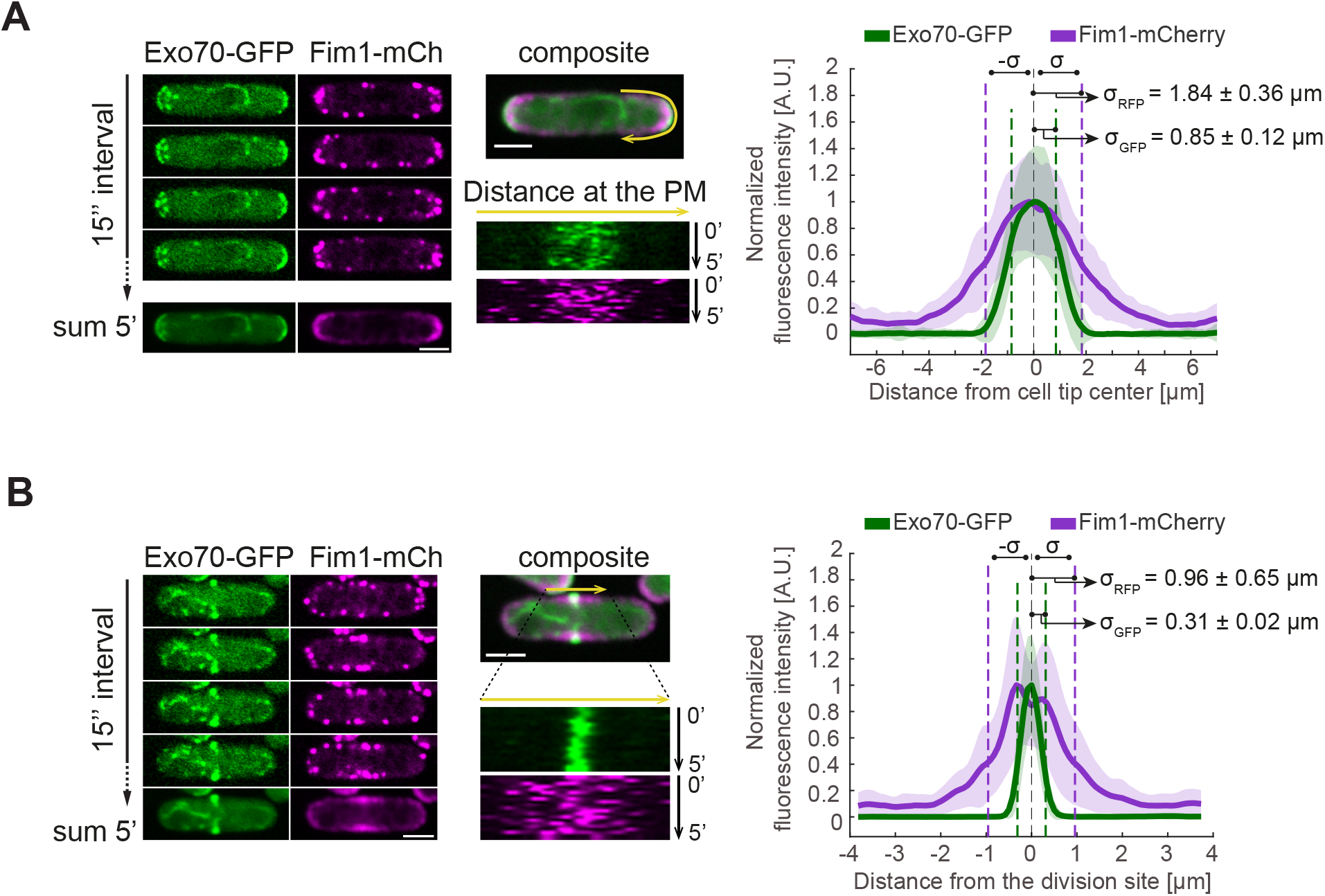
Distribution of exo- and endocytosis in fission yeast cells. **A-B.** Timelapse, sum projection and kymographs of the cell pole (A) and the early, pre-constriction division site (B) of cells expressing Exo70-GFP and Fim1-mCherry, marking exo- and endocytic sites, respectively. The graph shows the average distribution of these markers. Shaded areas show standard deviation.

**Figure S4.**
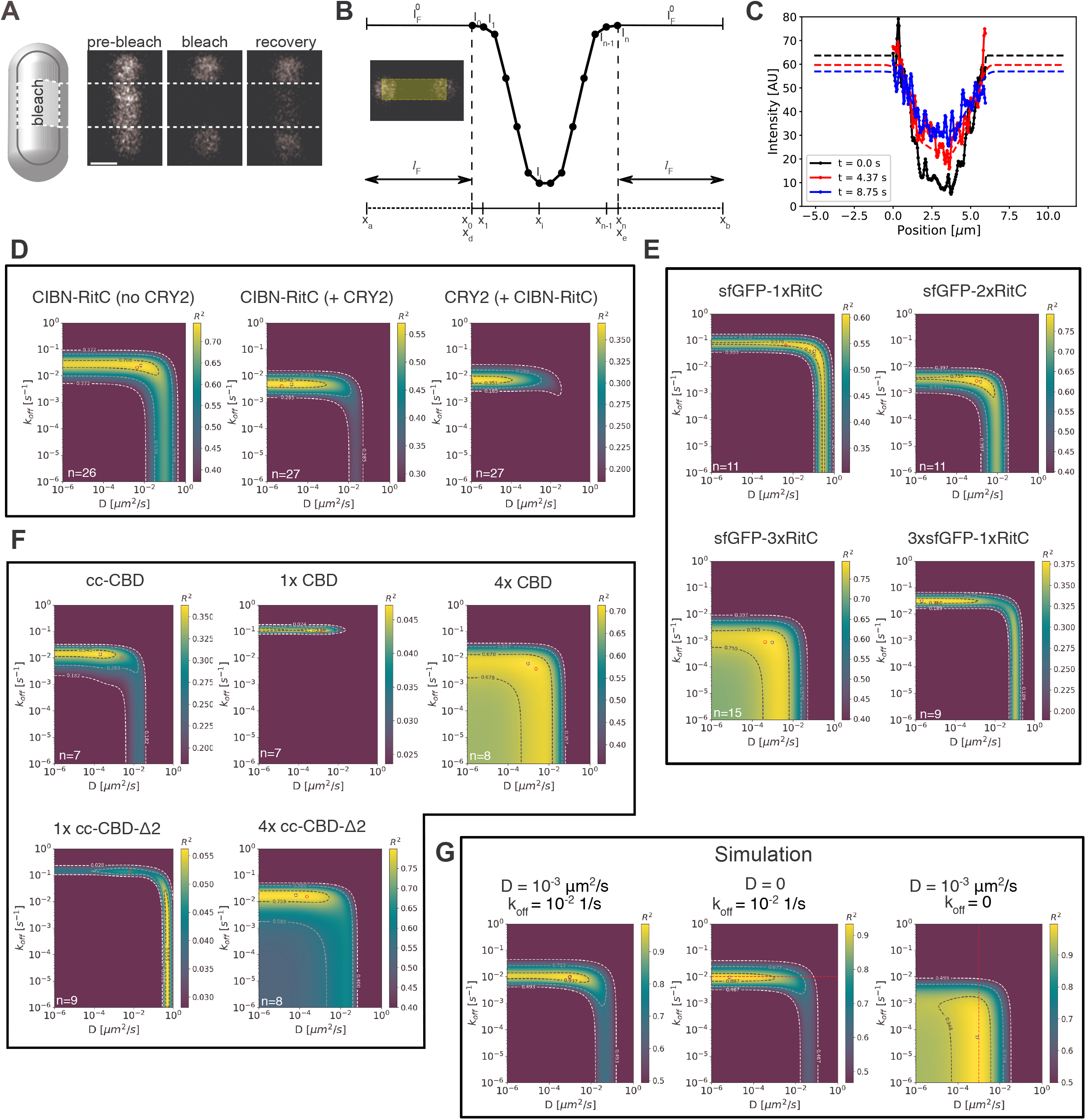
Fitting for diffusion coefficient and membrane dissociation rate with FRAP. **A.** A rectangular region along the cylindrical section on one side of the cell was photobleached and the recovery imaged over time. **B.** Schematic of 1D fluorescence recovery model for an experimental profile, extracted by vertical projection along a rectangular region shown in yellow, for the frame after photobleaching (*t* = 0, black circles). The boundaries between the experimentally-sampled middle region and the left and right flanking regions are indicated by dashed vertical lines. **C.** Example recovery curves. Single cell expressing 1xRitC-GFP imaged every 0.729 s and fit using the 1D model over 30 frames. Experimental data shown as lines with filled points while model fits shown as lines without points (*D* = 6.85 × 10^−3^ μm^2^/s, *k_off_* = 0.108 /*s* and *l_F_* as shown by the end points of the model curves). The black line of the model is initialized to go through the experimental data points at *t* = 0. **D-F.** Plots of averaged *R*^2^ calculated from FRAP fits for all protein constructs investigated as function of *D* and *k_off_*. Red circles indicate the median of the *D* and *k_off_* values for each protein from individual fits of the 1D model while blue dashed circles indicate the best fit using the average of individual *R*^2^ maps. The number of cells averaged in each plot is indicated by the value of *n*. **G.** Averaged *R*^2^ plots for simulated FRAP due to both diffusion and membrane binding/unbinding on the surface of a cylinder. Good agreement is found between the input parameters (red circle or dashed line) and the highest value of *R*^2^ (blue circle).

**Figure S5.**
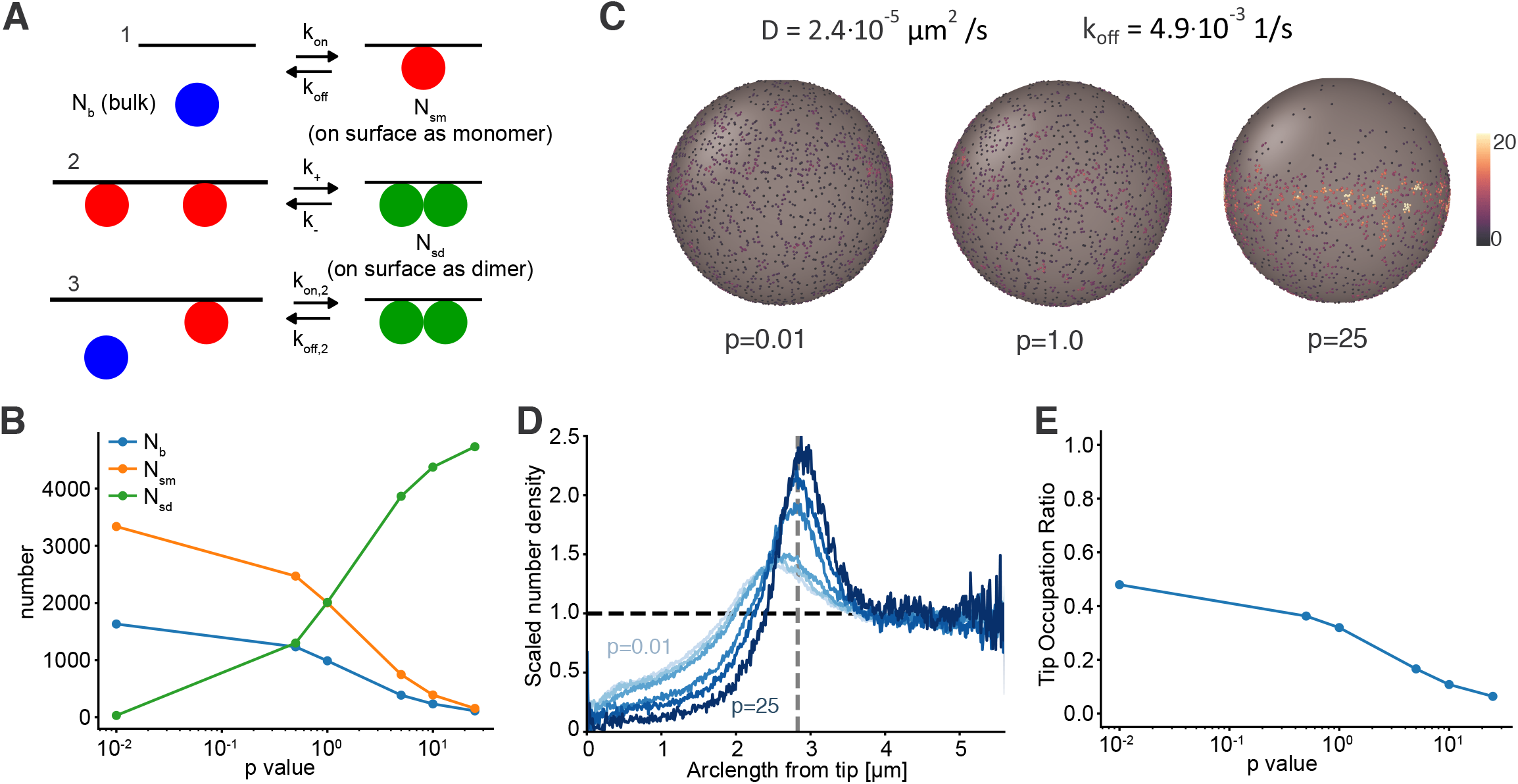
Modified model to include dimer formation on spherical surface, diffusing in the presence of membrane flows. **A.** Reactions used in dimerization simulations. The diffusion coefficient of dimers is the same as that of monomers. Reactions obey detailed balance at steady state in absence of endocytic or exocytic flow. **B.** Plots of the number of average particles at steady state as a function of parameter *p*. Simulations in this and following panels have 5000 particles and rate constants as described in supplementary text. **C.** Snapshots of spheres with increasing *p* value show that tip depletion and lateral concentration increase with increasing *p*. Color scale shows local density. **D.** Scaled number density of particles as a function of arclength show a similar trend where depletion and lateral peak increase with increasing *p*. **E.** Tip occupation ratio also decreases as a function of *p*.

**Figure S6.**
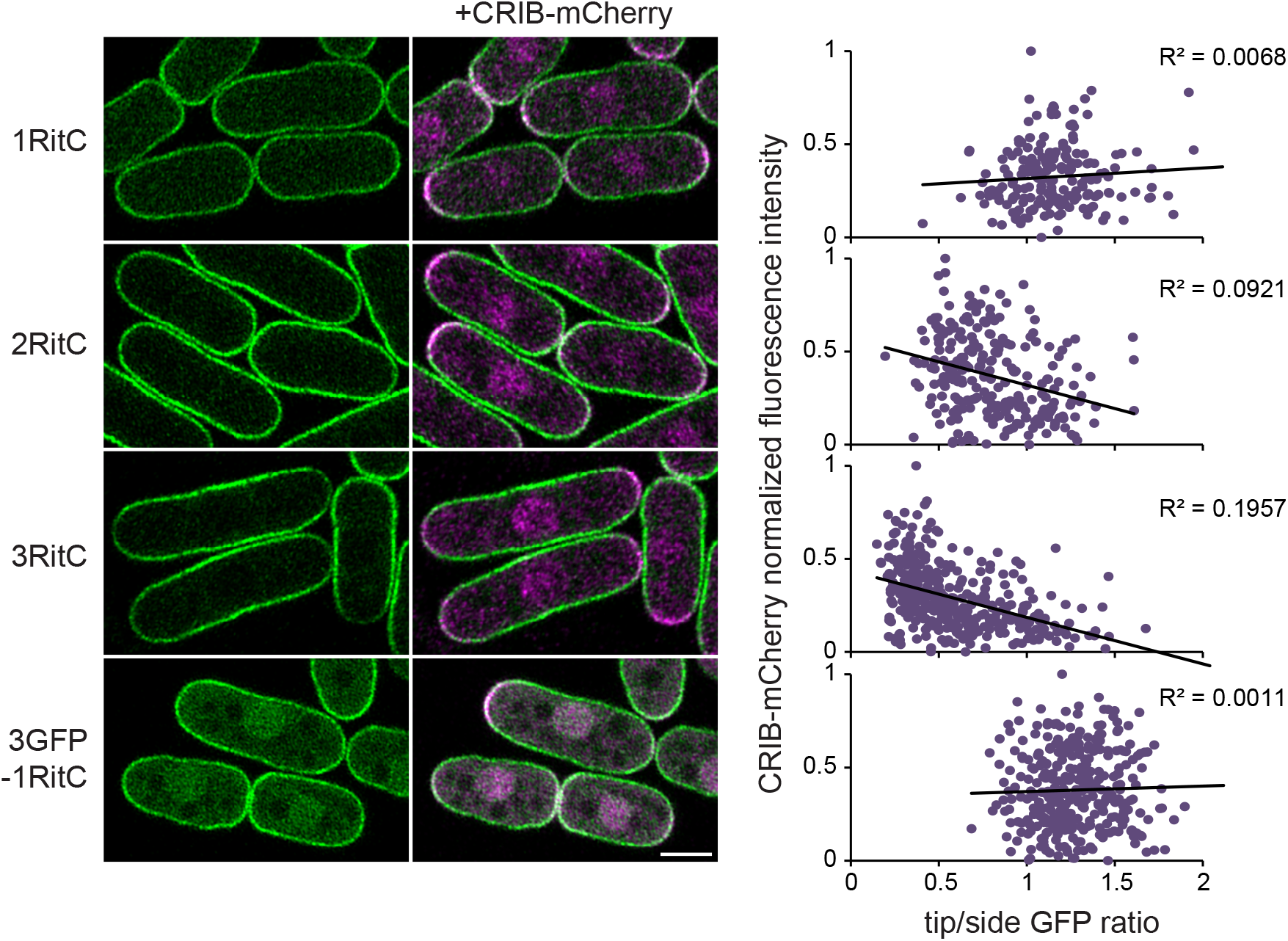
Membrane affinity-dependent depletion of membrane-associated RitC at sites of polarized secretion. Localization of indicated sfGFP-tagged RitC constructs in cells co-expressing CRIB-mCherry. The graphs on the right show correlation plots between the ratio of the tip to side GFP signal and the CRIB-mCherry tip intensity. Note the progressive shift towards the left of the data points upon RitC multimerization, as well as the more marked negative correlation.

**Figure S7.**
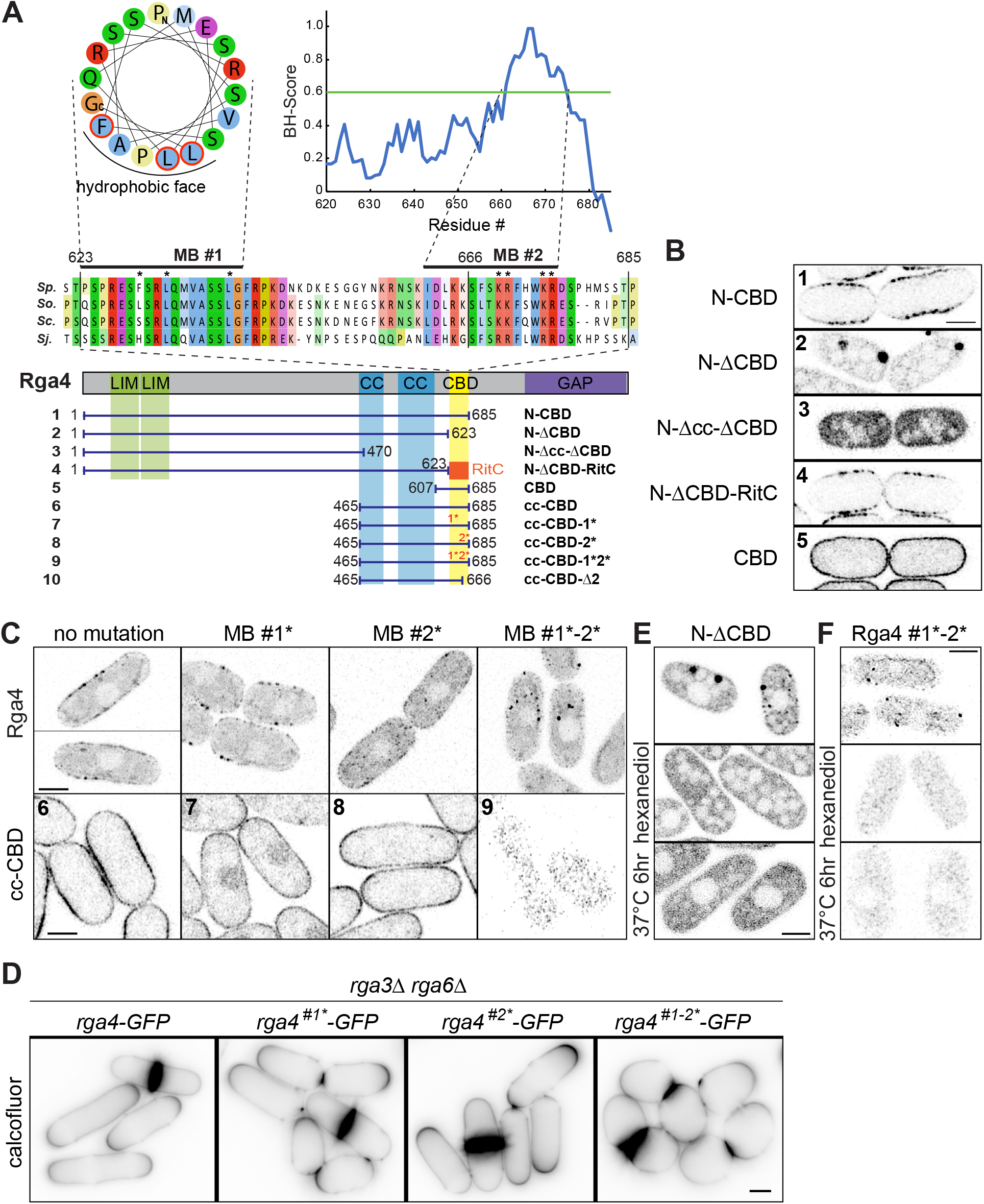
Structure-function analysis of the Cdc42 GAP Rga4 defining membrane-binding and oligomerization motifs. **A.** Scheme of Rga4 and truncation fragments. The sequence alignment shows the CBD across Schizosaccharomyces species (*S.p*. = *S. pombe; S.o*. = *S. octosporus; S.c*. = *S. cryophilus; S. j*. = *S. japonicus*). Membrane-binding (MB) motif 1 shows a weak amphipathic helix prediction, of which three hydrophobic residues (FLL, red circles and asterisks) were mutated to alanine. MB motif 2 was identified to have a high basic and hydrophobic (BH) scale, predictive of membrane association (*46*). Asterisks highlight four basic residues (KRKR) that were mutated to alanine. **B.** Localization of indicated Rga4 truncations, showing role of CBD in membrane localization and role of coiled coils (cc) in cluster formation. **C.** Localization of full-length Rga4 and the cc-CBD minimal Rga4 with or without mutations in MB #1 and #2. For full-length Rga4, max projection of 10 time points at 2s interval is shown. **D**. Shape of *rga3*Δ *rga6*Δ double mutant cells with indicated *rga4* alleles shown by calcofluor imaging. **E.** Dissolution of cytosolic condensates of Rga4 N-terminus lacking the CBD (N-ΔCBD) upon treatment with 10% 1,6-hexanediol or 37°C. **F.** Cytosolic condensates of full-length Rga4 with mutations in MB #1 and #2 are dissolved upon treatment with 5% 1,6-hexanediol or 37°C.

**Figure S8.**
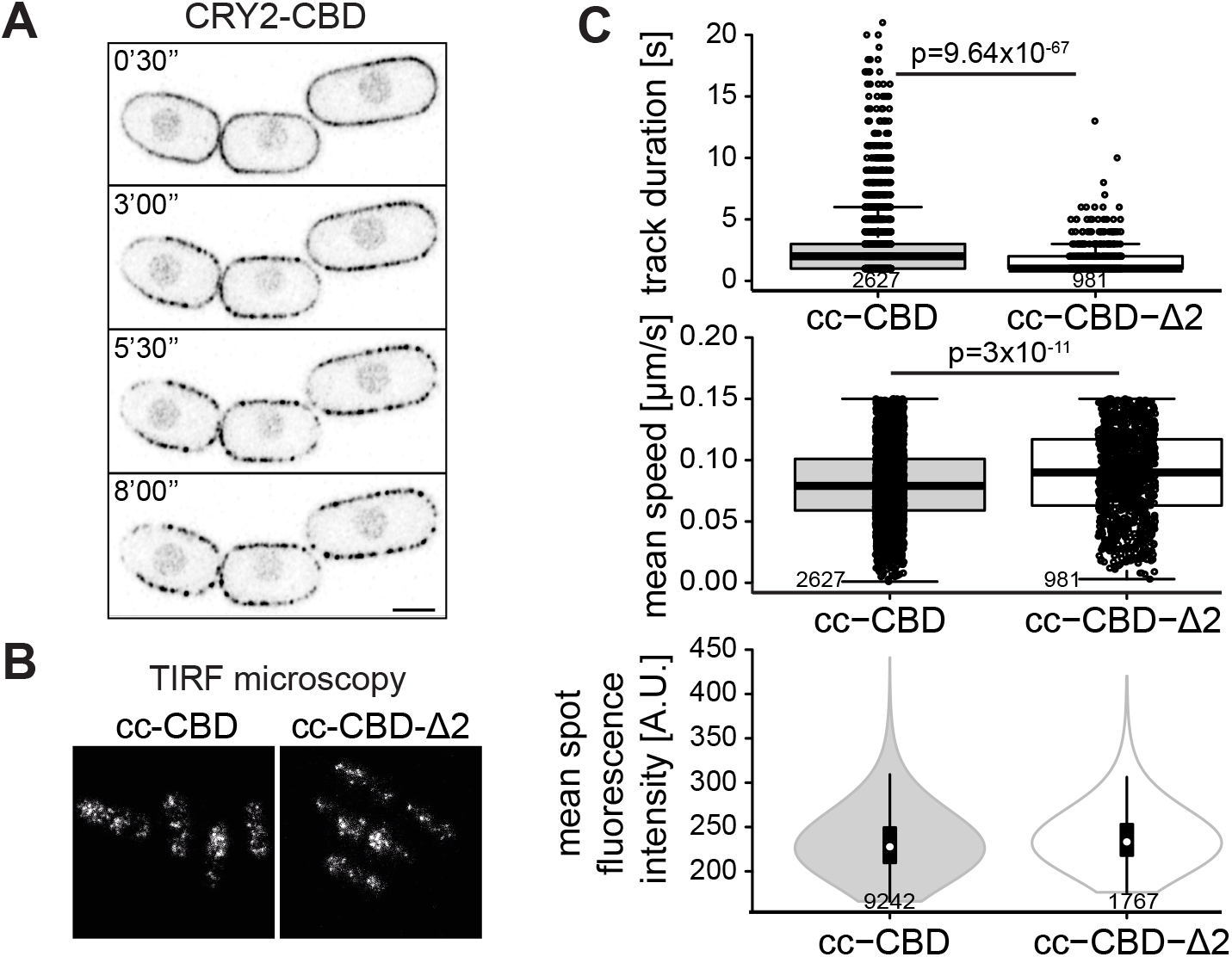
Further characterization of minimal Rga4 fragments. **A.** Localization of CRY2-CBD in cells grown in the dark and illuminated at time 0. **B.** Localization of cc-CBD and cc-CBD-Δ2 Rga4 fragments by TIRF-microscopy. See Movies 6-7 for time-lapse imaging and cluster tracking. **C.** Quantification of track duration, speed and fluorescence intensity for clusters of Rga4 fragment as in (B). The track duration of cc-CBD-Δ2 clusters was significantly reduced, while their speed was significantly increased, consistent with shorter membrane residence time and faster lateral diffusion. The track duration times and diffusion coefficient implied by the measurements (displacement per time step ≈ (4*D*t)^1/2^) are in the same range as the median *k_off_* and *D* of the FRAP fits in Fig S4F. The fluorescence intensity per cluster was not altered, suggesting unchanged oligomerization properties, as predicted from truncation of the membrane-binding region.

**Figure S9.**
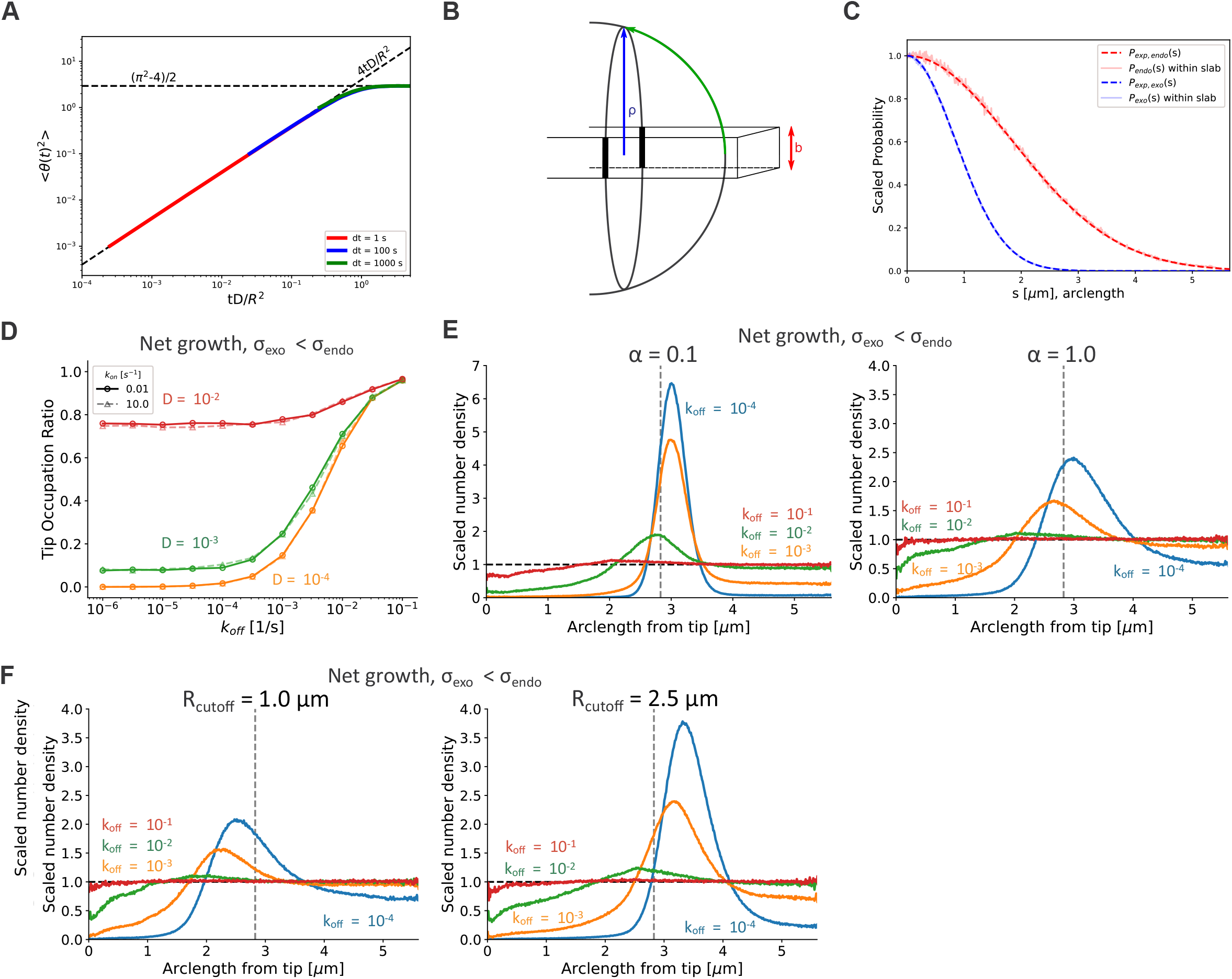
Model calibration and dependence on parameters *k_on_, α, R_cutoff_*. **A.** Test of implementation of diffusion on a sphere. Change in polar angle due to diffusion maintains the correct scaling of particle displacement over time, regardless of the timestep. Colored curves show results of averaged mean squared polar angle displacement (in radians squared) for particles diffusing on a sphere with *D* = 10^−3^ μm^2^/*s*, *R* = 2 μm. Dashed lines show asymptotic limits at short and long times from (*48*). The colored curves overlap, each starting from the smallest time, *t* = *dt*. **B.** Limited vertical resolution of microscope, *b*, leads to varying fractions of the cell surface being observed at a given arc length, *s*. **C.** Test of Equation (M4): if only events in a slab of height *b* are counted, sampling in 3D from *P*(*s*) recovers the experimentally observed probability distribution, *P_exp_(s*), for both the endocytosis and exocytosis profiles. **D.** Varying *k_on_* has no effect on the tip occupation ratio over the range of *D* and *k_off_* investigated, as expected. **E.** Parameter *α* (fraction of flowing membrane) affects tip depletion with lower (higher) values of *α* lead to enhanced (reduced) tip depletion and lateral peak. Other results in this paper used *α* = 0.5. **F.** Lower (higher) values of the cutoff range of exocytosis/endocytosis, *R_cutoff_*, leads to a reduction (increase) in tip depletion, along with reduction and leftward (increase and rightward) shift of the lateral peak. Other results in this paper used *R_cutoff_* = 2 μm.

### Movie legends

**Movie 1. Depletion of membrane-associated CRY2 at cell poles**

Time-lapse of CRY2-mCherry, CIBN-mTag-BFP2-RitC and CRIB-3GFP cells grown in the dark. Time 0 is the first timepoint after illumination. Images are at 30s interval. The UV channel (CIBN-mTagBFP2-RitC) was acquired every 4 timepoints. Depletion is observed at both poles of the three medial cells. Note that cells containing only CRY2-mCherry, which were used for normalization purposes, are also visible. Scale bar is 5μm.

**Movie 2. Depletion of membrane-associated CRY2 at division sites**

Time-lapse of CRY2-mCherry, CIBN-mTag-BFP2-RitC and CRIB-3GFP cells grown in the dark. Time 0 is the first timepoint after illumination. Images are at 30s interval. The UV channel (CIBN-mTagBFP2-RitC) was acquired every 4 timepoints. Depletion is observed initially at cell poles and then, as cells enter mitosis and prepare for cytokinesis at mid-cell, concomitant with CRIB-3GFP accumulation. Scale bar is 5μm.

**Movie 3. Simulation of membrane-associated protein depletion by membrane flows**

Simulation under static conditions (equal rates of membrane delivery by exo- and endocytosis), with k_off_ = 10^-5^/s and D = 10^-4^ μm^2^/s showing depletion from the zone of exocytosis on the top of the sphere.

**Movie 4. Simulation showing lack of membrane flow-dependent depletion for a protein with fast membrane unbinding rate**

Simulation under static conditions (equal rates of membrane delivery by exo- and endocytosis), with k_off_ = 10^-2^/s and D = 10^-4^ μm^2^/s showing absence of depletion from the zone of exocytosis on the top of the sphere.

**Movie 5. Localization of 3xRitC during the cell growth cycle**

Time-lapse of sfGFP-3xRitC and CRIB-3xmCherry in cells during mitotic growth. 3xRitC depletes from zones of growth labelled by CRIB at cell poles during interphase and repopulates the cell poles during mitosis when polarized cell growth stops. 3xRitC also transiently depletes from mid-cell, labelled by CRIB in pre-divisional cells until the initiation of septum invagination. White arrowheads show depletion zones. Scalebar is 5μm.

**Movie 6. Dynamics of Rga4 cc-CBD fragment in TIRF microscopy**

Tracking of cortical clusters of the Rga4 cc-CBD fragment in cells imaged by TIRF microscopy detected by TrackMate ImageJ plugin. Detected clusters are circles. All trajectories are shown.

**Movie 7. Dynamics of Rga4 cc-CBD-Δ2 fragment in TIRF**

Tracking of cortical clusters of the Rga4 cc-CBD–Δ2 fragment in cells imaged by TIRF microscopy detected by TrackMate ImageJ plugin. Detected clusters are circles. All trajectories are shown.

**Movie 8. Light-induced tube outgrowth in *rga3*Δ *rga4*Δ *rga6*Δ optoGAP cells**

Time-lapse of *rga3*Δ *rga4*Δ *rga6*Δ cells expressing CRIB-3GFP and optoGAP grown in the dark and illuminated with the blue light from t = 0. OptoGAP, shown in magenta, initially decorates the entire cell periphery, with CRIB absent from the cell cortex. The cells initiate polarized growth, with depletion of optoGAP, accumulation of CRIB in a restricted zone and outgrowth of a tube.

